# Antibody Fc receptor CD16a mediates natural killer cell activation via mechanotransduction of piconewton forces

**DOI:** 10.64898/2026.02.06.704366

**Authors:** Rong Ma, K. Christopher Garcia, Bianxiao Cui, Markus W. Covert

## Abstract

Natural Killer(NK) cells eliminate target cells through antibody-dependent cell-mediated cytotoxicity(ADCC), initiated by CD16a(FcγRIIIa) recognizing the Fc region of antibodies bound to the target cell surface. While the recognition is considered to be driven by CD16a-Fc binding avidity, it fails to explain why Fc multimers inhibit ADCC rather than trigger it. Here, we reveal that CD16a transduces piconewton forces, and acts as a mechanosensor to facilitate NK activation. We demonstrate that CD16a force and the actin foci formation associated with it are essential for the phosphorylation of mechanosensitive adaptor Cas-L and signaling adaptor LAT, reshaping NK cell cytoskeletal dynamics and signaling. Our findings show that NK activation is an intricate process that integrates both biochemical and biophysical information, and provides new mechanistic insight for immunoengineering.

---

Natural killer (NK) cells are lymphocytes in our innate immune system, protecting us from infections and cancer through their effector functions such as induced cytotoxicity and cytokine production. NK cell effector function is regulated by various receptor-ligand interactions at the cell-cell interface that are involved in target cell recognition. Following the discrimination of diseased cells from healthy cells, NK cells can kill target cells through a mechanism called antibody-dependent cell-mediated cytotoxicity (ADCC, Fig. 1A). ADCC is mediated by a Fc receptor, CD16a (FcγRIIIa), through a low-affinity interaction with the fragment crystallizable (Fc) region of an IgG1 antibody molecule that is bound to an antigen on the target cell surface (*1*). ADCC is also one of the major strategies harnessed in cancer immunotherapy through antibody treatments, such as trastuzumab and rituximab.

**Figure 1.**
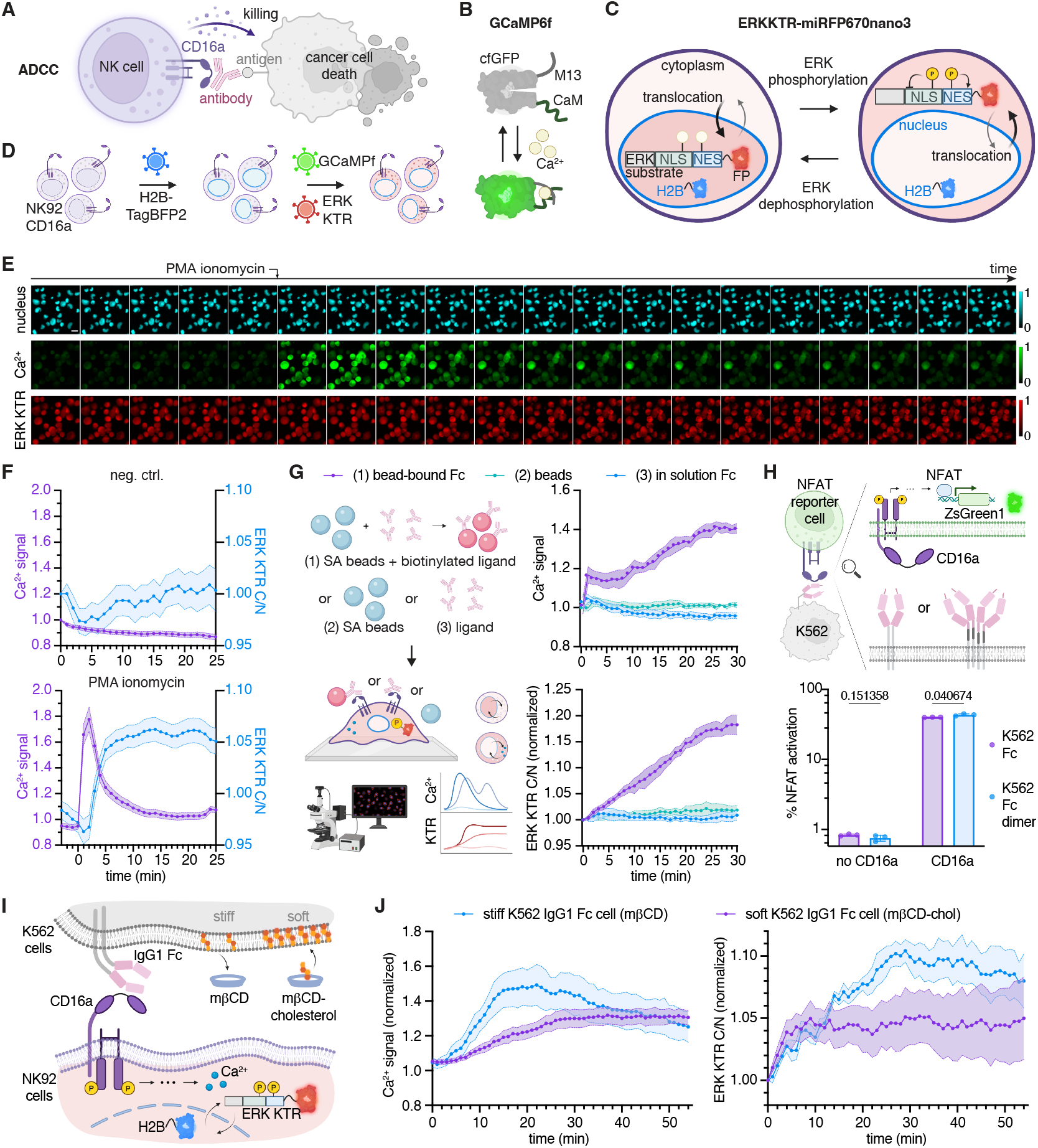
Mechanosensing contributes to potent NK cell activation by surface-bound IgG1 Fc. (A) Scheme illustrating ADCC mediated by CD16a in NK cells. (B) Scheme showing the GCaMP6f reporter used for monitoring Ca^2+^ flux. (C) Scheme showing the ERKKTR-miRFP670nano3 reporter used for monitoring ERK signaling. (D) Scheme illustrating the generation of NK92 CD16a reporter cell line by lentiviral transduction. (E) Representative images of reporter cells in H2B-TagBFP2 nucleus marker, Ca^2+^ signaling, and ERK signaling channels from t=-5 min to 15 min upon PMA ionomycin stimulation. Scale bar = 20 µm, interval = 1 min. Signaling activation of the entire population averaged from extracted single-cell traces. Data (mean±SEM) collected from 3 replicates with n=173 cells for negative control and n=233 cells for PMA ionomycin treatment. (G) Scheme and data showing that surface-bound Fc can trigger Ca^2+^ and ERK signaling potently compared to empty beads or Fc in solution in NK92 CD16a reporter cells. Data (mean±SEM) collected from 3 replicates, each replicate contains signaling traces of n=300-400 cells per condition. (H) Scheme and data show that two Fc expressed as one tethered dimer or four Fc expressed as 2 tethered dimers can activate CD16a with only ∼3% difference in NFAT activation (mean±SEM, 3 replicates). Statistical analysis was performed with unpaired two-tail t-test. (I) Scheme illustrating the experiment of co-incubating K562 IgG1 Fc target cells of different stiffness with NK92 CD16 reporter cells. (J) Data showing that stiffer K562 IgG1 Fc target cells can induce more potent Ca^2+^ flux and ERK phosphorylation in NK cells. Data (mean±SEM) collected from 3 replicates, each replicate contains single-cell traces of n∼200 cells per condition.

However, our mechanistic understanding of ADCC is very limited. One question pertains to how NK cells can distinguish antibody molecules that are bound to a target cell surface from the ones that are freely circulating. Given the low affinity of CD16a-Fc binding (∼ 5 µM), sufficient avidity is often considered to be the primary factor in addition to affinity for driving ADCC (*2*). Though CD16a-Fc binding is not a multivalent interaction (1:1 stoichiometry), it is currently hypothesized in the field that cell-bound antibodies would potentially induce CD16a clustering in NK cells, which initiates signaling (*3-5*). CD16a microcluster formation and the subsequent centralization leads to the assembly of NK immune synapse and enables ADCC effector function. Yet, this hypothesis is insufficient in explaining NK activation: engineered Fc multimers in solution can bind onto NK cells through multivalent binding and in principle bring several CD16a molecules together, evident by enhanced binding to cells using flow cytometry (*6*). Despite this enhanced avidity, these Fc multimers do not induce NK activation through ADCC without the Fab region facilitating immobilization to antigens on the target cell surface (*7*). On the contrary, ADCC can be inhibited by incubation with Fc multimers in solution, as was demonstrated with engineered Fc dimers, trimers, and hexamers (*6, 8-10*). Interestingly, a recent report found that CD16a exist as dimers 17 nm apart in resting NK cells before activation, leading to more mysteries of NK activation through CD16a (*5*). The current clustering hypotheses cannot fully explain the mechanism, underlining a major gap in our fundamental understanding of NK cell activation.

In contrast to the less understood CD16a-mediated NK activation in innate immunity, the mechanism of its counterpart, T-cell receptor (TCR)-mediated T cell activation in the adaptive immune system, is more explored (*11, 12*). Parallel to CD16a, TCR triggering is the primary signal to induce effector function in T cells, which also results in the release of cytotoxic granules to kill cancer cells. Moreover, while TCR directly recognizes antigens on cancer cell surfaces whereas CD16a does this indirectly, both receptors signal through the association with ITAM (immunoreceptor tyrosine-based activation motif) -bearing adaptors such as CD3ζ and FcγR (*13-16*). Growing evidence has revealed that T cells leverage mechanical forces through TCR in the process of recognizing antigens, initiating signaling, and exerting cytotoxic function (*17-20*). Though there have been a few speculations of NK cells being mechanosensitive (*21*), these studies have been limited to adhesion receptor LFA-1(*22*), pieozo-1 (*23*), and NKG2D (*24-26*). There was also one report that suggested CD16a could form catch or slip bonds with its nanobodies, however the study focused on measuring the binding kinetics of CD16a and its nanobody under flow in a cell-free system (*27*), lacking in-depth investigation of mechanical force transduction and biological significance.

Given the paralleled recognition and cytotoxic effector function of NK and T cells, we hypothesized that the primary immune activating receptor CD16a transmits mechanical forces and functions as a mechanosensor to dictate NK cell function.

To test this hypothesis, we used microscopy approaches to investigate cell-generated mechanical behavior through CD16a in both engineered NK92 CD16a reporter cells and primary human NK cells. Specifically, we used high-throughput live cell imaging to capture the signaling dynamics in NK cells and molecular tension fluorescence microscopy to visualize CD16a mechanical activity. We show that CD16a transmits defined pN forces against its antibody and natural ligand IgG1 Fc during NK activation. The CD16a force generation and transmission is associated with actin foci dynamics, and the foci formation is almost perfectly co-localized to total early total phosphorylation, as well as the phosphorylation of a key adaptor LAT (Linker of activation of T cells) and a mechanosensitive scaffold protein Cas-L. Our findings suggest a mechano-signaling mechanism in NK cells during early activation.

## Results

### Monitoring NK signaling dynamics with reporter cells

To study CD16a mechano-signaling, we first generated a NK92 reporter cell line to track the signaling dynamics in NK cells at single cell level upon activation. NK92 cells do not naturally express CD16a, making them ideal for this study. We engineered NK92 cells to express low-affinity CD16a (F158) and confirmed the expression with flow cytometry (Fig. S1A). To monitor NK signaling dynamics, we chose Ca^2+^ flux and ERK phosphorylation as NK activation markers. We used GCaMP6f as the Ca^2+^ sensor (Fig. 1B), which consists of a calmodulin, a circularly permuted green fluorescent protein (cfGFP), and a peptide M13. The binding to Ca^2+^ leads to conformational changes in GCaMP6f and the modulation of solvent access and pKa of the chromophore, resulting in fluorescence increase (*28*). We employed kinase translocation reporter (KTR) technology to obtain ERK phosphorylation dynamics (Fig. 1C). KTR is a fluorescent protein fused with a nuclear localization and exportation signal sequence (NLS, NES) and a specific kinase substrate for the kinase of interest. Depending on the target kinase activity, NLS and NES can get phosphorylated or dephosphorylated, leading to translocation into and out of the nucleus (*29*). This mechanism allows kinase activity information in individual cells to be obtained by extracting the cyto/nuclear fluorescence ratio from the translocated fluorescent protein, a far-red fluorescence protein miRFP670nano3. Therefore, we transduced the NK92 CD16a cells with lentivirus particles containing H2B-TagBFP2 to track cell nucleus, GCaMP6f to monitor Ca^2+^ flux, and ERKKTR-miRFP670nano3 to monitor ERK phosphorylation (Fig. 1D, Figure S1B). NK92 CD16a cells that expressed all constructs were isolated using FACS to establish the NK92 CD16a reporter cell line (Fig. S1C).

After generation of the NK92 CD16a reporter cell line, we tested whether these cells could report signaling dynamics by imaging them on positive and negative controls, which are poly-D-lysine (PDL) coated plates with or without PMA ionomycin stimulation. Cells were imaged with a 20× objective in a high-throughput manner so that the dynamic signaling of hundreds to thousands of cells can be monitored without biases. We adopted the previously reported image analysis pipeline CellTK for data extraction and analysis (Fig. S2, Video S1) (*30*). Representative images (Fig. 1E) and extracted activation traces of individual cells (Fig. S3) show that both GCaMP6f and ERKKTR functioned as expected, with the fluorescence of GFP increased after PMA and ionomycin addition, and subsequently the ERKKTR-miRFP670nano3 translocated from the nucleus to cytoplasm. By averaging the single cell traces we also obtained a population response, which showed that the calcium flux was triggered and peaked immediately, while ERK phosphorylation peaked at ∼ 5 min (Fig. 1F). We also confirmed that our reporter cell line is sensitive enough to respond to stimulations through CD16a at different antiCD16 coating concentrations (Fig. S4).

### NK cells are activated when CD16a interacts with immobilized, but not soluble, ligands

After establishing the NK92 CD16a reporter cell system, we first confirmed that NK signaling is triggered by surface-bound CD16a ligands (Fig. 1G). We immobilized biotinylated human IgG1 Fc onto streptavidin beads and monitored NK cell signaling as bead-bound Fc was introduced. Empty streptavidin beads and the same amount of soluble IgG1 Fc were used as negative controls. We observed that both Ca^2+^ and ERK phosphorylation were triggered by bead-bound IgG1 Fc but not by soluble Fc or empty beads (Fig. 1G, Fig. S5). We also confirmed this observation after Fc stimulation with an engineered Jurkat CD16a reporter cell line by measuring NFAT activity (Fig. S6). As this observation is often reasoned to be driven by avidity from multiple receptor-ligand binding events, we wondered if increasing CD16a-Fc binding valency would result in significantly better activation in both soluble form and at cell-cell interface. To test whether high valency of soluble Fc can trigger CD16a signaling, we expressed a previously reported IgG1 Fc hexamer and examined its ability to induce NK cell activation (*31*). We found that it indeed cannot induce Ca^2+^ or ERK phosphorylation at 1 µg/mL and 5 µg/mL (Fig. S7). To examine the effect of valency at cell-cell interface, we engineered K562 cell line to express tethered IgG1 Fc, so that they can be used to directly stimulate NK cells without the need of antigen presentation and antibody incubation such as CD20 and Rituximab. We found that tethered Fc dimer, despite potentially allowing two CD16a molecules to engage in proximity, offered only a very minor increase (∼3%) in NFAT activation compared to cells incubated with the Fc monomer (Fig. 1H and Fig. S8A). Overall, this data indicates that NK cells can sense the bound or unbound state of the antibody interacting with CD16a, suggesting that the physical resistance exerted by the bound antibodies might play a role in NK activation.

### Target cell stiffness influences NK cell activation

While induced CD16a clustering remains a possible contributor to NK activation through engaging surface-bound IgG1 Fc, the impact of Fc multivalency suggests that there could be other factors significantly driving this process. Therefore, we aimed to investigate whether the mechanism could be due to mechanosensing at the cell-cell interface. After confirming that K562 IgG1 Fc target cells can induce potent signaling when co-incubated with NK92 CD16a reporter cells (Fig. S8B), we tracked signaling dynamics when NK cells interact with target cells of different stiffness. We treated K562 IgG1 Fc cells with MβCD to remove cholesterol from cell membrane, which resulted in stiffer K562 IgG1 Fc cells in both membrane stiffness and cortical stiffness. We also treated K562 IgG1 Fc cells with MβCD-cholesterol to load more cholesterol into the cell membrane, resulting in softer target cells (*32*). The cholesterol removal and loading were confirmed by FilipinIII staining (Fig. S8C), and the stiff or soft K562 IgG1 Fc cells were subsequently introduced to NK92 CD16a reporter cells. We found that the stiffer cells can trigger Ca^2+^ flux and ERK phosphorylation more potently than softer ones (Fig. 1I, J, Video S2). These cell-cell interaction experiments indicate that NK cell activation may require a finely regulated mechanosensing machinery able to perceive the stiffness of target cells.

### CD16a transmits defined pN forces and acts as a mechanosensor in NK cells

We next sought to dissect the NK mechanosensing behavior through CD16a in a quantitative manner by coupling high-throughput imaging of cell signaling dynamics with molecular tension fluorescence microscopy (MTFM). DNA hairpin tension probes are a superior MTFM tool for receptor force studies, as they provide the highest spatiotemporal resolution as well as force sensitivity (*20, 33*). DNA hairpin tension probes use fluorophore-quencher pairs to report on the mechanical extension and unfolding of DNA hairpins under force (Fig. 2A, closed and open state of DNA hairpin tension probe). The hairpin is hybridized to a fluorescent (Cy3B) ligand strand on one arm and a quencher (BHQ2) anchor strand on the other arm, and subsequently immobilized on a glass substrate (Fig. S9A). In the absence of mechanical force, the hairpin is closed and thus the fluorescence is quenched; in the presence of a mechanical force applied to the ligand presented on top of the probe that is greater than the hairpin *F*_1/2_ (the force at equilibrium that leads to a 50% probability of unfolding), the hairpin mechanically unfolds and generates a fluorescent signal. This digital probe design is highly modular, which enables mapping forces above different magnitudes simply by using a hairpin with a different *F*_1/2_ (Fig. 2A, force sensing module). We presented both antiCD16 and IgG1 Fc to NK92 CD16a cells to evaluate their mechanical force profile (Fig. 2A, ligand presentation).

**Figure 2.**
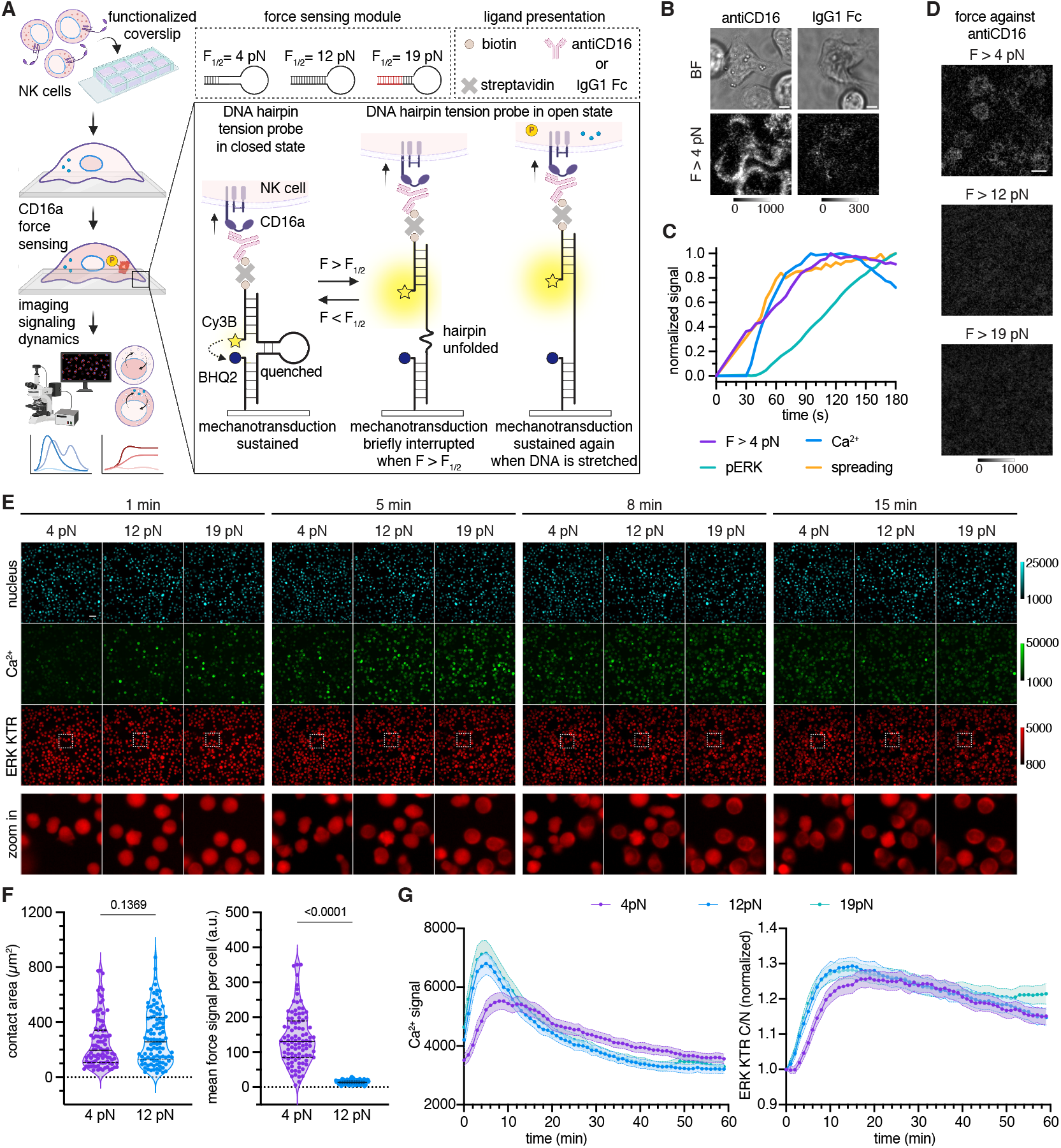
CD16a transmits defined pN forces and acts as a mechanosensor in NK92 cells. (A) Schematic showing the measurement of CD16a forces and signaling dynamics using DNA tension probes and NK92 CD16a reporter cells. (B) Representative 100× bright field and fluorescent images of CD16a forces against antiCD16 and IgG1 Fc. Scale bar = 5 µm. (C) Representative traces of sequential events during NK activation on antiCD16 DNA tension probe substrate in a representative NK92 CD16a reporter cell, including cell spreading, CD16a forces greater than 4 pN, Ca^2+^ triggering, and pERK signaling. Experiment was repeated three times. (D) Representative 20× fluorescent images showing NK92 CD16a reporter cells producing forces greater than 4 pN but less than 12 and 19 pN against antiCD16. Scale bar = 20 µm. (E) Representative fluorescent images of cell nucleus, Ca^2+^ flux, and ERKKTR translocation (with zoom in) during NK92 CD16a reporter cell activation on DNA tension probes presenting antiCD16 at different time points. Scale bar = 50 µm. (F) Contact area and mean force signal per cell of NK92 CD16a cells incubated on antiCD16 DNA tension probe substrates at t=10 min. Experiment was performed in three replicates, n = 97 and 102 cells. Statistical analysis was performed with unpaired two-tail t-test. Population response of NK92 CD16a reporter cells incubated on different DNA hairpin probes. Experiment was performed in three replicates, data averaged (mean±SEM) from single cell traces of Ca^2+^ triggering and ERK phosphorylation, n = 396 to 669 cells per condition.

We prepared substrates functionalized with 4, 12, and 19 pN DNA tension probes at the same density (Fig. S9B, C), and to ensure that we could pick up the potential transient and weak forces, we also used a previously reported lock oligonucleotide to amplify the tension signal over time by mechanically selective hybridization (Fig. S10A) (*19*). We imaged NK cells on the tension probe substrates presenting antiCD16 or IgG1 Fc, while substrates with no ligand served as a negative control (Fig. S10B). We found that NK cells were able to generate and transmit mechanical forces greater than 4 pN through CD16a during activation. The observed CD16a force against anti-CD16 exhibited a puncta pattern during cell spreading and was also located at the periphery of the cell (Fig. 2B, antiCD16). While CD16a forces greater than 4 pN against antiCD16 were consistently observed in NK92 cells, only a fraction of cells showed forces greater than 4 pN against IgG1 Fc, and in those that did, force signals were less abundant (Fig. 2B, IgG1 Fc, Video S3). Note this observation means that force transmission still exists through natural receptor-ligand binding, however it is weaker to unfold the 4 pN hairpins. Due to the difficulties in force visualization against the natural ligand Fc in NK92 CD16a reporter cells, we decided to focus on using antiCD16 to interrogate the mechanosensing mechanism through CD16a. Using CD16a antibody also has significant physiological relevance, as CD16a antibody single-chain variable-fragment (scFVs) are often used in bispecific or trispecific engager designs that aim to recruit NK cells for their cytotoxicity function (*34-36*). The We consistently observed that after the cell first contacted ligand presenting substrates, CD16a force generation accompanied cell spreading at a similar rate, followed by Ca^2+^ triggering and then ERK phosphorylation (Fig. 2C). The forces NK92 cells generated through individual CD16a against antiCD16 were mostly between 4 and 12 pN, as much less hairpin opening was observed with the 12 pN probe and 19 pN probe (Fig. 2D). Interestingly, with high throughput imaging of signaling dynamics, we noticed that though the DNA hairpin substrates at different F_1/2_ were almost identical chemically, the rate of NK activation at the population level was different. Representative images (Fig. 2E, Video S4) show that at 1 min, fewer cells on 4 pN probe exhibited Ca^2+^ triggering than cells seeded on 12 and 19 pN probes. At 5 min, though cells on all substrates showed Ca^2+^ signal increase, cells on 12 and 19 pN substrates had a stronger response accompanied with ERKKTR translocation. By 8 min, cells on all substrates showed similar level of Ca^2+^, while there were still more cells on 12 and 19 pN substrates with ERKKTR translocation. Later at 15 min, the Ca^2+^ and ERK activity looks identical for all three substrates. Quantitative analysis of cell spreading area and force signal (Fig. 2F), as well as the extracted signaling traces (Fig. 2G) indicates that the hairpin opening slightly negatively impacted signaling, evident by a slower onset and smaller peak in Ca^2+^ signaling, as well as a slower response in ERK phosphorylation. We suspect this observation was due to brief interruption of mechanotransduction with 4 pN hairpin unfolding (Fig. 2A, mechanotransduction) and further supports the significant role of CD16a mechanosensing in NK activation. Overall, we combined live-cell imaging of receptor forces and NK intracellular signaling to quantify the mechanical forces at the piconewton scale at which NK are activated by CD16a.

### Actin dynamics and foci formation contributes to CD16a mechanics and NK activation

As we identified CD16a as a mechanically active receptor, we next aimed to elucidate cellular input that could contribute to the CD16a mechanical activity. We chose small-molecule inhibitors CK666 and latrunculin B (lat B) to pretreat NK92 CD16a reporter cells for 20 min to inhibit Arp2/3 in actin branching as well as actin polymerization prior to incubating cells on PDL or antiCD16 coated wells (Fig. 3A). Compared to cells on antiCD16 without treatment or with only DMSO, both CK666 and lat B abolished ERK phosphorylation in NK activation through CD16a (Fig. 3B). This inhibition of activation was also confirmed in NFAT signaling using Jurkat CD16a NFAT reporter cells (Fig. S11).

**Figure 3.**
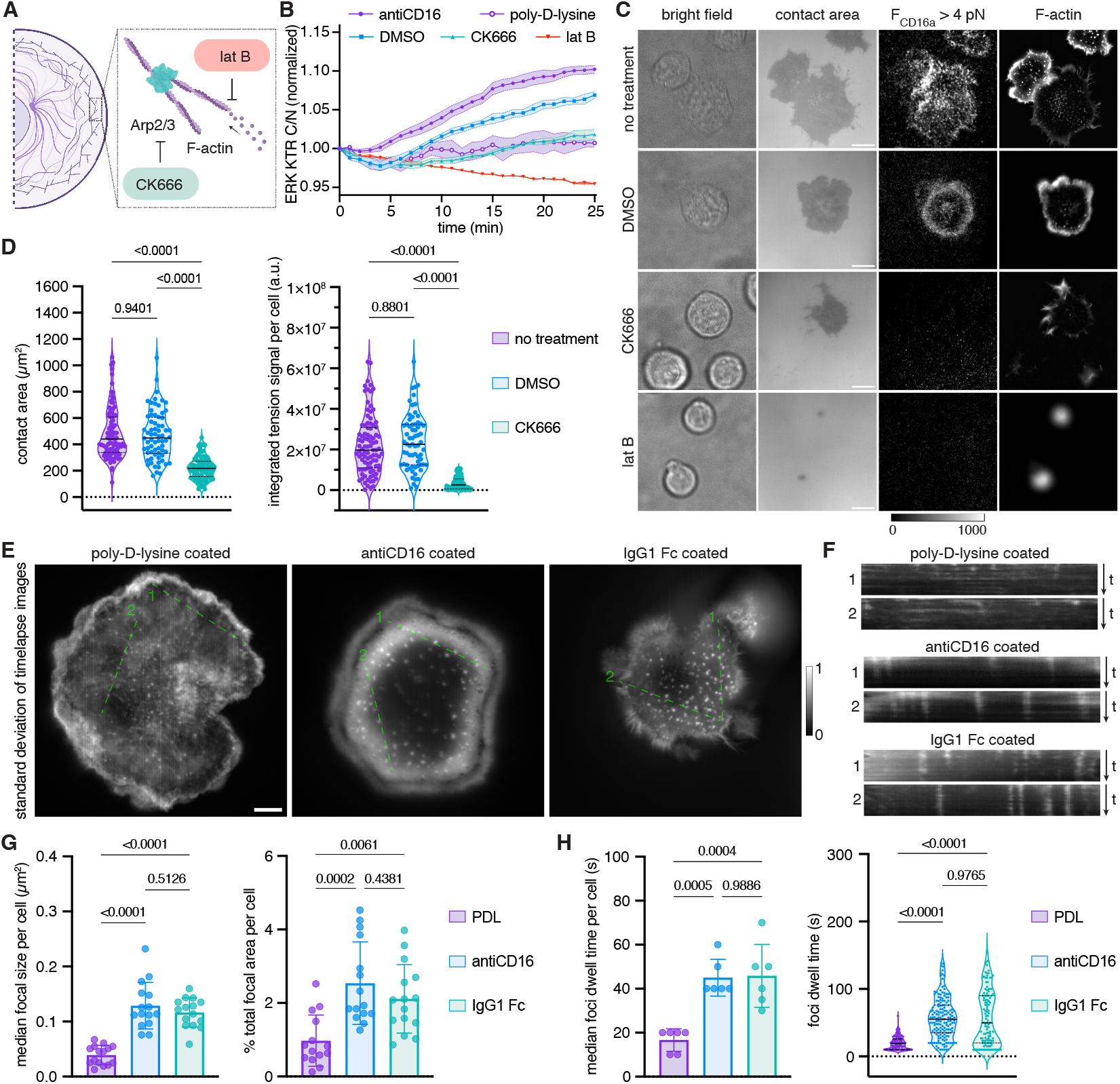
Actin dynamics affect the forces and are involved in NK activation through CD16a. (A) Schematic showing the inhibition of specific cytoskeletal activities. (B) Extracted ERK traces show the NK activation is inhibited with small molecule inhibitor lat B and CK666. Data shows mean±SEM from 3 replicates, and each condition contains traces of n ∼ 200 cells per replicate. (C) Representative bright field, contact area, tension and actin images of NK92 CD16a cells treated with DMSO, CK666, lat B, or no treatment. Scale bar = 5 µm. (D) Quantitative analysis on contact area and forces. Plot show data collected from cells in three replicates, n = 95, 68, 100 cells. Solid line indicates the median and dashed line indicates the quartiles. Statistical analysis was performed with one-way ANOVA and Tukey’s multiple comparison. (E) Standard Deviation images of F-actin timelapse in representative NK92 CD16a lifeact cells from t = 5 min to 10 min on activating and non-activating substrates, interval = 10 s. Scale bar = 5 µm. (F) Kymographs of ROIs in representative cells marked in (E). (G) Plots show the median focal size per cell and the percentage of total focal area occupied of the cell contact area at t = 5 min. Data was analyzed from representative cells from three replicates. Mean±SD, n = 15 cells per condition, and foci counts range from 23-290 per cell. Statistical analysis was performed with one-way ANOVA and Tukey’s multiple comparison. (H) Dwell time of actin foci for cells incubated on PDL, antiCD16 and IgG1 Fc substrates from t = 5 min to 10 min, mean±SD. Dwell time was obtained by particle tracking analysis and statistical analysis was performed with one-way ANOVA and Tukey’s multiple comparison. Data was collected from n = 6 representative cells and foci counts = 544, 460, 378 in 3 replicates.

We then examined the effect of inhibiting cytoskeletal activities on NK cell spreading and CD16a force transmission using 4 pN DNA tension probes presenting antiCD16. The inhibition of Arp2/3 by CK666 reduced NK spreading area and dramatically decreased observed forces through CD16a (Fig. 3C, D). Cells treated with lat B did not spread or produce CD16a forces and thus we did not include it in quantitative analysis. To obtain more mechanistic insight, we engineered NK92 CD16a cells to express a F-actin marker lifeact-mNeonGreen for these experiments (Fig. S12). Though some cell spreading was maintained upon CK666 treatment, we found that the formation of actin foci was eliminated (Fig. 3C, F-actin). Hence, we suspected that actin foci play a role in NK activation.

To further investigate actin foci formation, we followed their dynamics on activating ligand coated glass compared to non-activating PDL coated glass. The standard deviation of timelapse images (Fig. 3E, Video S5) of NK92 CD16a lifeact cells showed the most dynamic F-actin changes during image acquisition, from which we found that actin foci were more pronounced on activating substrates than PDL substrate. Kymographs of actin foci (ROI 1 and 2 marked, green dash in Fig. 3E) showed that actin foci on activating ligands were more stable (Fig. 3F). Quantitatively, on antiCD16 and IgG1 Fc subtrates, there was no significant difference in size of actin foci and % area actin foci occupied of total contact area per cell, while both substrates yielded significantly larger actin foci than cells spreading on PDL (Fig. 3G, Fig. S13). Particle tracking analysis by TrackMate (Fig. 3H, Video S6) showed that the median dwell time of individual actin foci was ∼ 40 - 50 s on antiCD16 and IgG1 Fc substrates, significantly longer than that on PDL substrate. In summary, we found that actin cytoskeletal activities contribute significantly to NK cell activation, in part due to the formation of relatively larger actin foci that have a longer dwell time when engaging activating ligands on the tested planar surfaces.

### CD16a force-associated actin foci are phosphorylation sites for the adaptor proteins Cas-L/LAT

Based on these observations, we next sought to explore the connection between actin foci, CD16a force, and NK activation. We imaged accumulated CD16a forces using NK92 CD16a cells expressing liveAct-mNeonGreen as they spread and exert force on 4 pN DNA hairpin probes presenting antiCD16 with the lock oligo (Video S7). We found that the puncta pattern in CD16a forces resembled the actin foci formation. To find their association patterns, we summed the images from the first 200 s in the F-actin channel as well as the tension channel (Fig. 4A). Images were marked with ROI 1 (a linescan) and ROI 2 (5 small squares) in the F-actin channel and in the tension channel. Figure 4B shows images of F-actin and tension of the same region from 20 to 60 s, with 2 more ROIs (ROI 3 and 4, arrows) labeled on the images. In the ROI 1 linescan on the summed image, we found that actin foci colocalized with sites of accumulated CD16a force signal (Fig. 4C). As observed in ROI 3 and 4 in Fig. 4B, we noticed that actin foci were assembled first prior to CD16a force transmission exceeding 4 pN. Based on this, we tracked the intensity of the randomly picked positions on sum images (ROI 2) and observed that actin assembly frequently took place 10 to 20 s prior to the transmission of CD16a forces that were greater than 4 pN (Fig. 4D, Cyan arrows). Similarly, with a bit of lag, actin disassembly was often associated with no new CD16a force transmission, leading to a plateau in the force plot (Fig. 4D, black arrows).

**Fig. 4.**
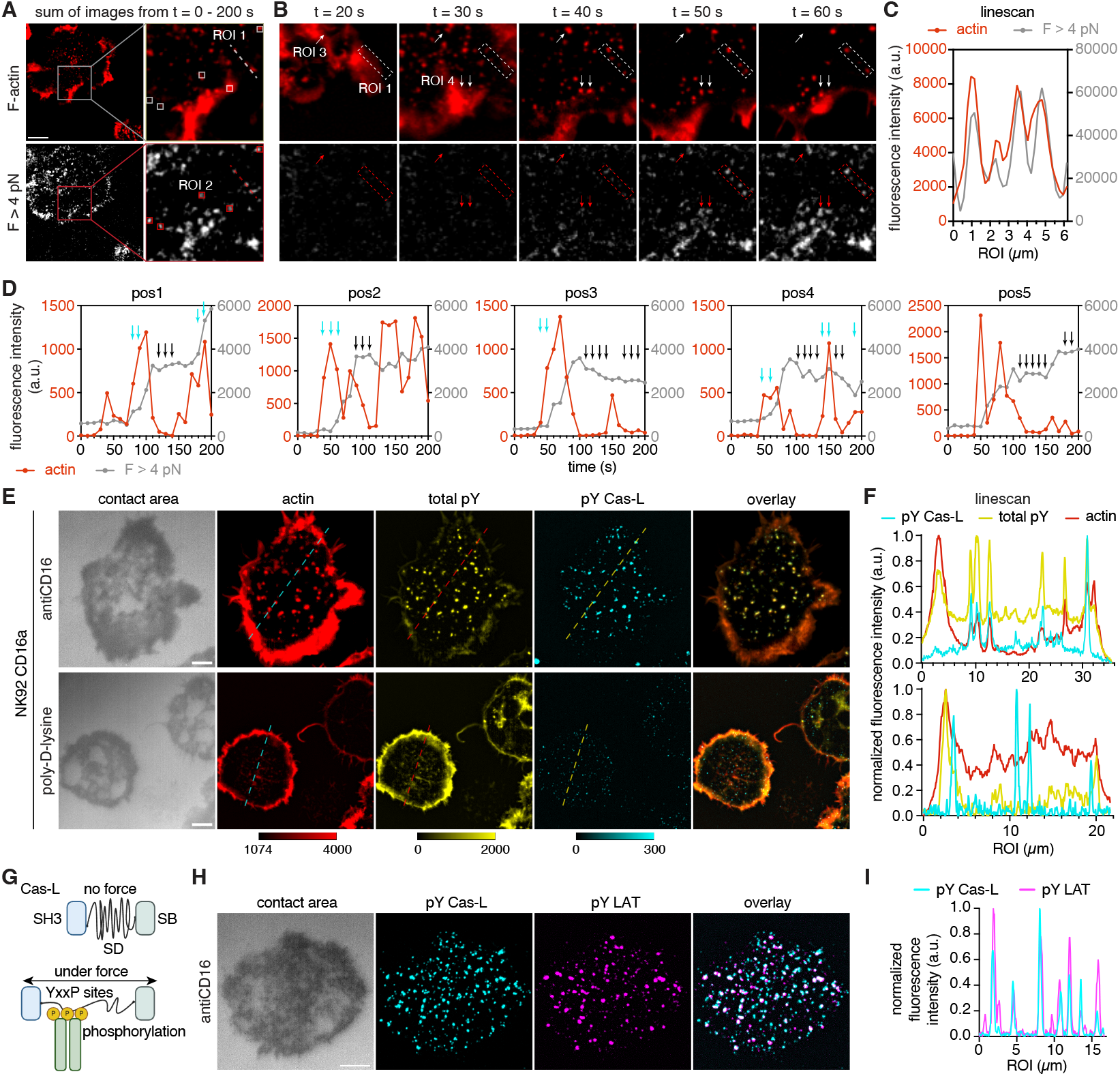
CD16a force is associated with actin foci and sites of signaling adaptor phosphorylation. (A) Sum of timelapse F-actin and accumulated force images from t = 0 - 200 s from a representative cell. Column on the right is a zoom-in view of the box with two ROIs, ROI 1 is a linescan and ROI 2 are five 4×4 pixel boxes. Scale bar = 5 µm. (B) Timelapse F-actin and accumulated tension images of the zoom-in view with two more ROIs marked by arrows. (C) Linescan of ROI 1 in sum images showing the colocalization between F-actin and tension channels. (D) Time course plots of F-actin and accumulated tension signal for ROI 2. Cyan arrows mark the actin foci assembly followed by accumulated force signal increase, while black arrows mark the disassembly of actin foci followed by accumulated force signal plateauing. (E) Representative immunofluorescence staining images of actin, total pY, and pY Cas-L in NK92 CD16a cells incubated on antiCD16 or PDL substrates at t = 10 min. Scale bar = 5 µm. (F) Linescan of the dashed line ROI marked in (E) showing the colocalization of actin foci, total pY and pY Cas-L. Fluorescence intensity in each channel was normalized. (G) Scheme illustrating mechanosensing mechanism by Cas-L. (H) Representative immunofluorescence staining images of pY Cas-L and pY LAT in NK92 CD16a cells incubated on antiCD16 substrate at t = 10 min. Scale bar = 5 µm. (I) Linescan of the dashed line ROI marked in (H) showing the colocalization of pY Cas-L and pY LAT. Fluorescence intensity in each channel was normalized.

While it was exciting to establish the connection between actin foci and CD16a forces, we were curious about the signaling function of these sites of force-foci association. After incubation on substrates coated with either antiCD16 or PDL (neg. ctrl), NK92 CD16a cells were fixed and stained for F-actin and total phosphorylation (Fig. 4E) We found that during early NK cell activation, actin foci were co-localized with sites of total phosphorylation (Fig. 4F). Since actin foci are associated to sites of CD16a force transmission and total phosphorylation, we next wondered if any specific intracellular protein could be involved in the CD16a mechanotransduction. P130Cas is one of the mechanosensing adaptors involved in cell adhesion, and it has been found that the phosphorylation YxxP sites in the substrate domain are only exposed if the protein is stretched (*37*). It is shown that the most abundant member of Cas family proteins in lymphocytes is Cas-L (Fig. 4G), which can affect T cell signaling and activation (*38-40*). We stained for the phosphorylation of Cas-L, and found that it also co-localized to actin foci and total pY (Fig. 4E, F), with the pY Cas-L and total pY Pearson’s correlation coefficient = 0.81 ± 0.06 (mean±SEM). To further establish the mechanotransduction mechanism between CD16a force, actin foci, and phosphorylation on Cas-L, we sought to determine if any signaling adaptor could be involved. Thus, we stained for the phosphorylation of LAT, which is a key adaptor protein in immune cell signaling and known to be responsive upon Fc receptor stimulation (*41*). We found that the phosphorylation of LAT also co-localized with phosphorylation of Cas-L (Pearson’s correlation coefficient = 0.52 ± 0.03, mean±SEM), especially to the larger patches (Fig. 4H, I). Therefore, we conclude that the signal propagation during NK activation is mediated by a mechanism involving mechanosensing through CD16a and mechanotransduction through Cas-L.

### CD16a transmits higher magnitude of forces for mechanosensing in primary NK cells

Last, we confirmed the mechanical activity of CD16a in primary NK cells. We purified primary human NK cells from peripheral mononuclear cells (PBMCs) and imaged the cells using substrates functionalized with DNA hairpin tension probes (Fig. 5A). Primary NK cells can indeed exert defined forces through CD16a to IgG1 Fc compared to negative control (Fig S14), and we consistently observed accumulated forces against IgG1 Fc (Video S8), in between 4 pN to 19 pN (Fig. 5B). Surprisingly, CD16a forces against antiCD16 in primary NK cells were higher than that in NK92 cell line (Fig. 5C, Video S9). Quantitative analysis also showed that cell spreading was affected by different hairpin mechanical threshold, with accumulated forces greater than 19 pN against antiCD16 (Fig. 5D). Timelapse imaging of real-time forces show that as soon as the cell made an initial contact with ligands, force was immediately transmitted through CD16a with a puncta pattern (Fig. 5E, 10 to 30 s), and then expanded to the periphery of the cell as the cell spread (70 - 120 s), consistent with what we see in NK92 CD16a cells (Video S7).

**Fig. 5.**
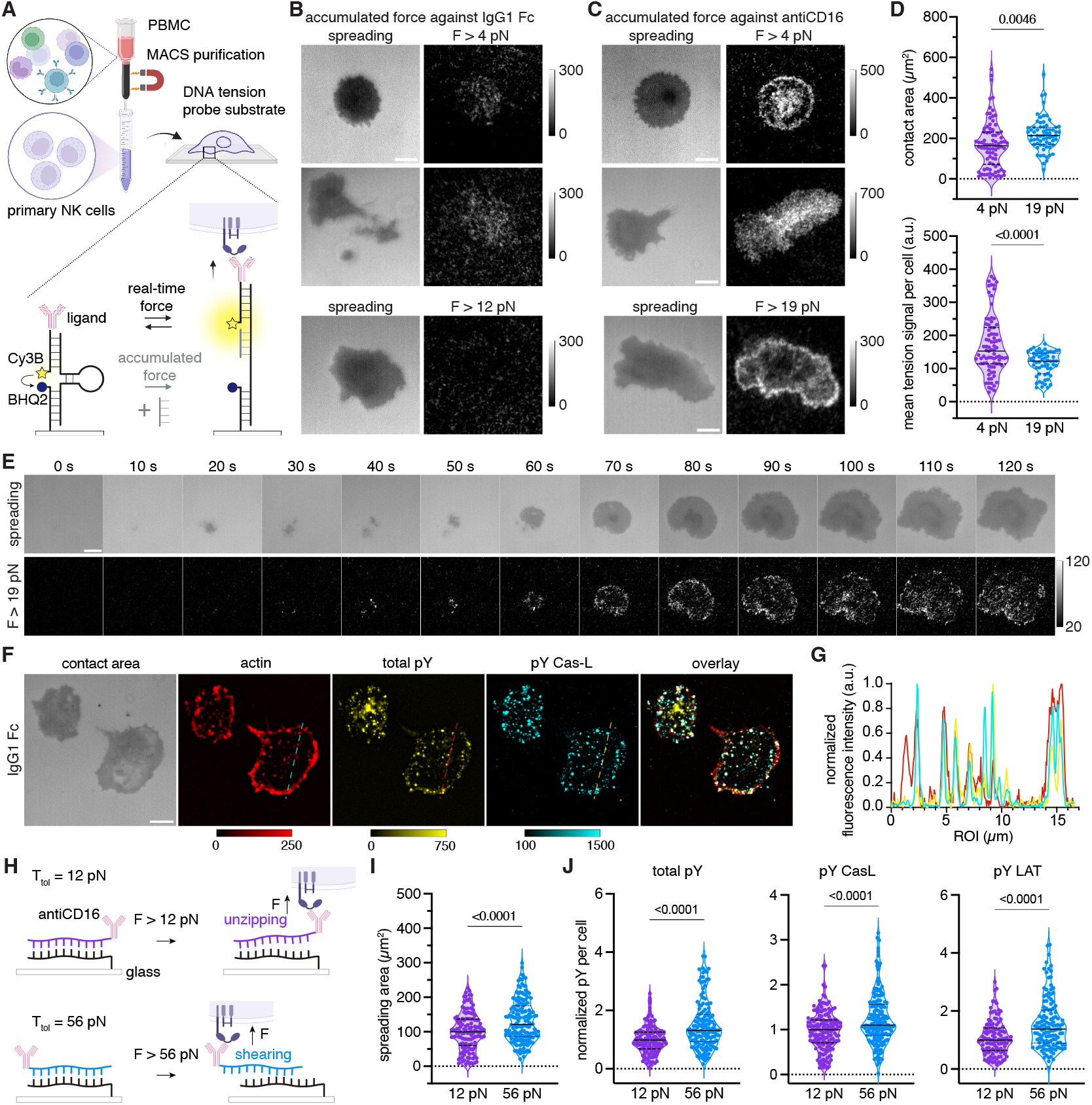
CD16a transmits pN forces and senses different mechanical environments in primary NK cells. (A) Schematic shows the MACS purification of primary NK cells from PBMC and force measurements on DNA tension probe substrates in real-time or accumulated manner (shown in grey). (B) Representative images showing CD16a forces generated by primary human NK cells against IgG1 Fc were greater than 4 pN but smaller than 12 pN. Scale bar = 5 µm. (C) Representative images showing CD16a forces generated by primary human NK cells against antiCD16 were greater than 19 pN. Scale bar = 5 µm. (D) Quantitative analysis shows the contact area and accumulated CD16a force against antiCD16 in primary NK cells. Data were collected from three replicates, solid line represents median and dashed lines represent quartiles. (E) Timelapse images of a representative primary NK cell spreading and exerting real-time force greater than 19 pN against antiCD16. Scale bar = 5 µm. (F) Representative immunofluorescence staining images of actin, total pY, and pY Cas-L in primary NK cells incubated on IgG1 Fc substrates at t = 10 min. Scale bar = 5 µm. (G) Linescan of the dashed line ROI marked in (F) showing the colocalization of actin foci, total pY and pY Cas-L. Fluorescence intensity in each channel was normalized. (H) Schematic showing TGTs as tools to manipulate the force threshold allowed for mechanotransduction. As soon as the TGT ruptures, the mechanotransduction is terminated. (I) Spreading area and (J) total, Cas-L, and LAT phosphorylation in primary NK cells on TGT presenting antiCD16 at t = 5 min. Mean±SD, data collected from 3 replicates, n ∼ 160 cells per condition, solid line indicates median and dashed line indicates quartiles.

We also confirmed that actin foci still co-localized to total phosphorylation and phosphorylation of the mechanosensing Cas-L protein in primary NK cells (Fig. 5F, G), which possibly facilitated force transmission together. We then verified CD16a mechanosensing mechanism utilizing the tension gauge tethers (TGT) to present antiCD16, since TGTs can provide a higher force threshold which can withstand the greater than 19 pN forces that CD16a transmits to antiCD16 in primary NK cells. TGTs are chemically identical DNA duplexes that can be mechanically separated in unzipping or shearing geometry, leading to duplex rupture at either 12 or 56 pN and subsequent termination of mechanotransduction through CD16a (Fig. 5H). We found that primary NK cells exhibited larger contact area (Fig. 5I) and higher total phosphorylation, Cas-L phosphorylation, and LAT phosphorylation on 56 pN TGT rather than 12 pN (Fig. 5J). This is likely a result of more sustained mechanotransduction on 56 pN TGT, as CD16a forces in primary NK cells are above 12 and even 19 pN against antiCD16 and thus can easily rupture the 12 pN TGT and terminate CD16a mechanotransduction. The difference of phosphorylation on 56 and 12 pN TGTs further illustrates the mechanosensing ability of primary NK cells, which echoes the observations in NK92 CD16a reporter cells, though the force magnitude and mechanosensing thresholds differed. (Fig. 2G). These differences could be dependent on the intrinsic biophysical properties and cytoskeletal dynamics of NK92 cells and primary NK cells.

## Discussion

Surface-bound ligands can induce potent immune responses in both the innate and adaptive immune system. Examples include activation through TCR in T cells, B cell receptor (BCR) in B cells, and Fc receptor in NK cells. One widely accepted activation mechanism is that the binding to ligands can induce receptor clustering, which increases the local density of ITAMs and thereby increases the rate of phosphorylation on kinases and adaptors for signaling (*42*) (*13*). However, clustering cannot explain activation sufficiently. For T cell activation, a recent study showed that TCR engaging a single pMHC was enough for triggering activation signal (*43*). In the case of Fc receptor mediated activation, Fc multimers lacking antigen binding abilities but with enhanced binding to Fc receptors can actually inhibit antibody mediated effector functions including cytotoxicity (ADCC) and phagocytosis (ADCP), and thus are used as therapeutic strategies for autoimmune conditions. Taken together, these observations suggested that there could be other driving forces behind activation through the Fc receptor CD16a.

NK cells and CD16a’s counterparts, T cells and TCR triggering has been investigated extensively at both the molecular and cellular level, with significant findings emerging from a variety of perspectives, for instance, models of kinetic proofreading, mechanosensing, phase separation, and size exclusion. (*44-47*). These models greatly contributed to explaining the activation mechanism through TCR. However, to date NK activation lacks such extensive investigation, especially with newer technologies. As an important process that drives ADCC, which is harnessed in many therapeutical antibodies, the insufficient mechanistic understanding greatly limits our ability to design novel immunotherapies that recruit NK cells.

This work fills the gap by interrogating NK activation process via primary activating receptor CD16a from a biophysical perspective. We found that CD16a is a mechanically active receptor, capable of exerting forces greater than 4 pN to IgG1 Fc during NK activation. Marrying signaling reporters and MTFM, we were able to track the force events and dynamic signaling at the same time. As mechanical engagement of receptor ligand propagated, the Ca^2+^ flux was triggered and followed by sustained ERK signaling. One important observation was that when cells first encountered ligands on 4 pN hairpin tension probe, it took longer for them to trigger. We reason that the cells can unfold the 4 pN hairpins and thus don’t sense the reaction from immobilized ligand until the unfolded hairpin is fully stretched, which led to a slower activation after exerting a force through CD16a at the population level.

Given that small molecule inhibition of cytoskeleton impaired both CD16a forces and NK activation, we suspected that actin foci, a pattern specific to activating substrates driven by Arp2/3, are involved in NK mechanosensing ability through CD16a. Actin foci have been observed in immune cells incubated on 2-D planar substrates. Studies at cell-substrate interface have revealed that actin foci are leveraged by B cells to extract antigens (*48*), and that they are important in T cell activation and lytic immune synapse formation (*49, 50*). It is also reported that actin foci co-localize to TCR microclusters during T cell activation (*51*), and that the Arp2/3 facilitated fast F-actin turnover rate that likely contributes to the fast response from T cells (*50, 52*). These studies prompted us to investigate the connection between cytoskeleton, CD16a force, and NK mechanosensing. Consistent with the studies in T cells and B cells, we also found relatively large actin foci formed on both antiCD16 and IgG1 Fc substrates with an average dwell time around 50 s, which we suspect to be likely involved in mechanotransduction and signal propagation. Indeed, we were able to identify actin foci as part of the sites of where CD16a mechanotransduction took place, evident by the colocalization to the sites of total phosphorylation including Cas-L. Due to the nature of how Cas-L has to be stretched on both ends to reveal the phosphorylation sites, our finding elucidated a potential mechanosensing network. Though we were able to observe that force transmission through CD16a follows actin foci assembly, we currently have not yet determined whether the CD16a force transmission occurs before or after Cas-L phosphorylation. Thus, an important remaining question is whether the phosphorylation on Cas-L scaffold protein is a necessary condition for relaying CD16a force or if it is a response from CD16a mechanosensing and NK activation signal propagation? Answering this question is not only helpful in understanding NK activation, but also can provide insight in other immune cell triggering mechanisms.

Following our findings in the NK92 cell line, we further validated our results in primary human NK cells. CD16a indeed can transmit forces, however the force was greater than 19 pN to antiCD16 and only greater than 4pN to IgG1 Fc. Consistent with our force measurements, we also observed differences in activation level by phosphorylation at a higher threshold force of 56 pN compared to 12 pN, supporting our finding of NK mechanosensing through CD16a. However, different force magnitude raises another interesting question: what contributes to the force generation and sensing differences between primary NK cells and NK92 cell line? While CD16a polymorphism (158 F/V) could contribute to the differences in force profile due to their differential ability to bind to IgG1 Fc (*53*), the binding strength between NK92 and primary NK cells with the pan antiCD16 clone 3G8 should not affected. If immune receptor mechanosensing machinery is a pure intrinsic property of the molecular interaction interface, we should see similar force profile in NK92 and primary NK cells.

Mechanistically, while there are paralleled processes in T cells through TCR and NK cells through CD16a, there are also plenty of differences in these two systems. First, CD16a is regulated by ADAM17, which results in CD16a shedding upon NK activation (*54*). Whether ADAM17 could regulate NK mechanosensing abilities and potentially lead to NK cell fate of expansion or exhaustion is unknown. Second, unlike TCRs that are polyclonal, CD16a has limited polymorphism, which could contribute to differential mechanosensing abilities. For example, one of the most known variants is CD16a F158V, which has a higher affinity to IgG1 Fc and found in a smaller percent of the population (*55*). This difference in affinity also raised a third critical question in this comparison: in T cell mechanosensing, the low affinity TCR-pMHC interactions often exhibit a distinct catch-bond behavior, which is thought to minimize the interference by non-specific antigens and amplify signals specific to cognate antigens (*18, 56, 57*). As a low affinity interaction, do CD16a-Fc exhibit catch-bond behavior? Fundamentally, this comparison leads to the question of is catch-bond a phenomenon that can be explained on structural basis at molecular level, or is it a phenomenon initiated by cells with an intricate machinery to prolong the receptor-ligand ligation? This point also reinstates the question we raised when comparing the force magnitude in NK92 cell line and primary NK cells: despite the same molecular interaction, the mechanics can still be different when live cells exert forces themselves. Fourth, Fc glycosylation is known to affect the binding between Fc and Fc receptors, and engineered Fc variants can achieve enhanced ADCC and ADCP by afucosylation (*58, 59*). How does effector mechanosensing change along with engineered Fc? Despite the improved effector function, excessive inflammation can be induced by afucosylated Fc. Additionally, IgG glycosylation pattern changes in aging and diseases (*60*), which makes understanding how Fc glycosylation state regulates mechanosensing through CD16a another important direction. Fifth, the Fc region of antibodies have varied affinity to different Fc receptors, and thus can engage and recruit differential pathways and immune cells to orchestrate complicated immune response. Are other Fc receptors mechanically active? Sixth, NK cells are known to possess two major subsets: CD56^bright^, which is more involved in producing cytokines and immunomodulation; and CD56^dim^, which is more involved in exerting cytotoxic effector function. Stemming from the biochemical marker of CD56, on the biophysical side would we also see corresponding mechanical phenotypes of NK cells? Would stronger ability to exert forces correlate with their killing abilities, and are the killing events associated with certain pulling forces at the NK synapse? These are all important questions in understanding the similarities and differences of cytotoxic effector function in T cells and NK cells, which will contribute to a deeper understanding of immune signaling at cell-cell interface.

In addition to the mechanistic insights and questions, our findings also imply a potential approach for antibody-based immunotherapy that is based on mechanosensing rather than affinity-based engineering. Though at the moment it is unclear whether CD16a-Fc forms a catch bond, we can envision that Fc variants which can facilitate better mechanotransduction would potentially be more potent in recruiting NK cells for ADCC. Additionally, in the TCR engineering field, it has been shown that TCR catch-bond engineering can generate superior TCR clones with high potency but low off-target properties (*56*). We can foresee that receptor mechanics-guided engineering could potentially be powerful in designing new immunotherapies.

One of the limitations of this study is that our measurements were performed on glass coverslips. Though there have been speculations about the true form of actin foci at 3-D cell-cell junction, there is no doubt that 2-D planar glass substrate offers a great way to isolate and study the contribution from cytoskeleton. As foci structures and CD16a microclusters were previously observed on activating supported lipid bilayers (SLB) (*61, 62*), a promising approach would be to interrogate the actin foci dynamics with respect to receptor mechanics on SLBs as well. We suspect the actin foci could be protrusions (*63*), where CD16a on the cell membrane can quickly push and pull with the fast actin turnover under Arp2/3. Ideally, super-resolution imaging would be helpful to resolve the local mechanotransduction network through CD16a. It would also be highly desirable to track the dynamics of CD16a microcluster formation, force generation, and centralization movement with Cas-L phosphorylation on SLBs, and thus resolve the kinetics of signalosome assembly in the context of force requirement and generation (*64*). At the cellular level, this study focuses on population response in Ca^2+^ and ERK phosphorylation, it would also be desirable to include more reporters involved in different stages of activation and resolve step-wise signaling kinetics at single cell level with respect to mechanical engagement. Additionally, we did not interrogate any potential oscillatory Ca^2+^ or ERK signaling at single cell level, and it would be highly interesting to understand if there is any mechanical interplay. Another limitation in this study is that we could not resolve the contribution of mechanotransduction and clustering. The field has long established the significance of avidity in antibody mediated effector function without exploring a potential mechanosensing mechanism (*2*). While we report on CD16a mechanical activity, it is inherently difficult to fully isolate receptor mechanics from naturally formed receptor clusters. Engineered Fc multimers have demonstrated that though inducing multivalent interaction, they inhibit ADCC or ADCP processes. Perhaps an ultimate test would require the fabrication of DNA origami to limit the number of CD16a-Fc engagement in a given area, while allowing or terminating mechanical forces. On that note, studies have to carefully design the size of DNA origami platforms from the energy perspective (*65*), as they are quite rigid and might be able to provide reaction from CD16a pulling, since the CD16a-Fc forces are only ∼ 4 pN. Perhaps ultimately forces and clustering are always intertwined, and they collectively contribute to a cell’s ability for antigen discrimination: individual receptors within the clusters could potentially withstand higher forces and facilitate signalosome assembly. Moreover phase separation might also be contributing to forming such signaling condensates that can do more “heavy lifting” (*45*).

In conclusion, we report on a mechanosensing mechanism through the Fc receptor CD16a during NK cell activation. At a fundamental level, this research establishes CD16 as a mechanosensor, akin to TCR and BCR (*66*), which expands our understanding of Fc receptor signaling, NK cell activation, and broadens our knowledge in immunobiophysics. We expect our findings to inspire more mechanistic reevaluation of other members of FcγR family. In the future, we hope our findings can inspire new ideas in antibody engineering for immunotherapy, beyond the current affinity-driven approaches, and enable more effective harnessing of ADCC.

## Funding

R.M. acknowledges the support from Stanford Science Fellow program and Michelson Medical Research Foundation for the support through the 2021 Human Vaccines Project Michelson Prizes for Human Immunology and Vaccine Research. R.M. acknowledges the feedback from Dr. Khalid Salaita, Dr. Brian Evavold and Dr. Brendan Deal on this work. K.C.G. acknowledges the support from National Institutes of Health through NIH-AI103867 and Howard Hughes Medical Institute. All authors thank the Life Science Editors for editing services.

## Author contributions

Conceptualization: R.M.

Methodology: R.M.

Investigation: R.M.

Supervision: R.M., M.C., K.C.G., B.C.

Writing – original draft: R.M.

Writing – review & editing: R.M., M.C., B.C., K.C.G.

## Diversity, equity, ethics, and inclusion

Receptor tension not political tension!

## Competing interests

Authors declare that they have no competing interests.

## Data and materials availability

The data in this study are available upon request. The materials generated in this study are available upon request, but we may require a payment for shipping and/or a completed materials transfer agreement. The raw and extracted imaging data files in this study are available upon request. The analysis pipeline was adapted from previous publication (doi.org/10.1016/j.jbc.2023.104599), the pipeline documentation is publicly available at https://github.com/sjeknic/CellTK, https://celltk.readthedocs.io/en/latest/.

## Supplementary Materials

Materials and Methods

Figs. S1 to S14

Movies S1 to S9

## Notes

### Competing Interest Statement

The authors have declared no competing interest.

## References

1. E. Vivier, J. A. Nunes, F. Vely, Natural killer cell signaling pathways. Science 306, 1517–1519 (2004).

2. S. C. Oostindie, G. A. Lazar, J. Schuurman, P. Parren, Avidity in antibody effector functions and biotherapeutic drug design. Nat Rev Drug Discov 21, 715–735 (2022).

3. M. Dorsch et al., Quantitative analysis of human NK cell reactivity using latex beads coated with defined amounts of antibodies. Eur J Immunol 50, 656–665 (2020).

4. S. W. de Taeye et al., FcgammaR Binding and ADCC Activity of Human IgG Allotypes. Front Immunol 11, 740 (2020).

5. P. Ross et al., Spatial localization of CD16a at the human NK cell ADCC lytic synapse. bioRxiv, (2024).

6. X. Zhang et al., A recombinant human IgG1 Fc multimer designed to mimic the active fraction of IVIG in autoimmunity. JCI Insight 4, (2019).

7. E. A. Fitzpatrick, J. Wang, S. E. Strome, Engineering of Fc Multimers as a Protein Therapy for Autoimmune Disease. Front Immunol 11, 496 (2020).

8. R. Spirig et al., rIgG1 Fc Hexamer Inhibits Antibody-Mediated Autoimmune Disease via Effects on Complement and FcgammaRs. J Immunol 200, 2542–2553 (2018).

9. D. F. Ortiz et al., Elucidating the interplay between IgG-Fc valency and FcgammaR activation for the design of immune complex inhibitors. Sci Transl Med 8, 365ra158 (2016).

10. L. Tradtrantip, C. M. Felix, R. Spirig, A. B. Morelli, A. S. Verkman, Recombinant IgG1 Fc hexamers block cytotoxicity and pathological changes in experimental in vitro and rat models of neuromyelitis optica. Neuropharmacology 133, 345–353 (2018).

11. G. Gaud, R. Lesourne, P. E. Love, Regulatory mechanisms in T cell receptor signalling. Nat Rev Immunol 18, 485–497 (2018).

12. K. Shah, A. Al-Haidari, J. M. Sun, J. U. Kazi, T cell receptor (TCR) signaling in health and disease. Signal Transduct Tar 6, (2021).

13. A. H. Courtney, W. L. Lo, A. Weiss, TCR Signaling: Mechanisms of Initiation and Propagation. Trends Biochem Sci 43, 108–123 (2018).

14. L. L. Lanier, Up on the tightrope: natural killer cell activation and inhibition. Nat Immunol 9, 495–502 (2008).

15. A. Blázquez-Moreno et al., Transmembrane features governing Fc receptor CD16A assembly with CD16A signaling adaptor molecules. P Natl Acad Sci USA 114, E5645–E5654 (2017).

16. H. Medjouel Khlifi, S. Guia, E. Vivier, E. Narni-Mancinelli, Role of the ITAM-Bearing Receptors Expressed by Natural Killer Cells in Cancer. Front Immunol 13, 898745 (2022).

17. R. Basu et al., Cytotoxic T Cells Use Mechanical Force to Potentiate Target Cell Killing. Cell 165, 100–110 (2016).

18. M. A. Faust, V. J. Rase, T. J. Lamb, B. D. Evavold, What’s the Catch? The Significance of Catch Bonds in T Cell Activation. J Immunol 211, 333–342 (2023).

19. R. Ma et al., DNA probes that store mechanical information reveal transient piconewton forces applied by T cells. P Natl Acad Sci USA 116, 16949–16954 (2019).

20. Y. Liu et al., DNA-based nanoparticle tension sensors reveal that T-cell receptors transmit defined pN forces to their antigens for enhanced fidelity. P Natl Acad Sci USA 113, 5610–5615 (2016).

21. G. Santoni et al., Mechanosensation and Mechanotransduction in Natural Killer Cells. Front Immunol 12, (2021).

22. D. Friedman et al., Natural killer cell immune synapse formation and cytotoxicity are controlled by tension of the target interface. J Cell Sci 134, (2021).

23. A. K. Yanamandra et al., PIEZO1-mediated mechanosensing governs NK-cell killing efficiency and infiltration in three-dimensional matrices. Eur J Immunol 54, e2350693 (2024).

24. L. Mordechay et al., Mechanical Regulation of the Cytotoxic Activity of Natural Killer Cells. ACS Biomater Sci Eng 7, 122–132 (2021).

25. G. Le Saux et al., Nanoscale Mechanosensing of Natural Killer Cells is Revealed by Antigen-Functionalized Nanowires. Adv Mater 31, e1805954 (2019).

26. J. Fan et al., NKG2D discriminates diverse ligands through selectively mechano-regulated ligand conformational changes. EMBO J 41, e107739 (2022).

27. C. González et al., Nanobody-CD16 Catch Bond Reveals NK Cell Mechanosensitivity. Biophys J 116, 1516–1526 (2019).

28. T. W. Chen et al., Ultrasensitive fluorescent proteins for imaging neuronal activity. Nature 499, 295–300 (2013).

29. G. Maryu et al., Live-cell Imaging with Genetically Encoded Protein Kinase Activity Reporters. Cell Struct Funct 43, 61–74 (2018).

30. S. Jeknic, T. Kudo, J. J. Song, M. W. Covert, An optimized reporter of the transcription factor hypoxia-inducible factor 1alpha reveals complex HIF-1alpha activation dynamics in single cells. J Biol Chem 299, 104599 (2023).

31. H. Liu, A. Saxena, S. S. Sidhu, D. Wu, Fc Engineering for Developing Therapeutic Bispecific Antibodies and Novel Scaffolds. Front Immunol 8, 38 (2017).

32. K. Lei et al., Cancer-cell stiffening via cholesterol depletion enhances adoptive T-cell immunotherapy. Nat Biomed Eng 5, 1411–1425 (2021).

33. Y. Zhang, C. Ge, C. Zhu, K. Salaita, DNA-based digital tension probes reveal integrin forces during early cell adhesion. Nat Commun 5, 5167 (2014).

34. A. El-Khoueiry et al., First-in-Human Phase I Study of a CD16A Bispecific Innate Cell Engager, AFM24, Targeting EGFR-Expressing Solid Tumors. Clin Cancer Res 31, 1257–1267 (2025).

35. J. Harwardt et al., Generation of a symmetrical trispecific NK cell engager based on a two-in-one antibody. Front Immunol 14, 1170042 (2023).

36. S. Kakiuchi-Kiyota et al., A BCMA/CD16A bispecific innate cell engager for the treatment of multiple myeloma. Leukemia 36, 1006–1014 (2022).

37. Y. Sawada et al., Force sensing by mechanical extension of the Src family kinase substrate p130Cas. Cell 127, 1015–1026 (2006).

38. M. Minegishi et al., Structure and function of Cas-L, a 105-kD Crk-associated substrate-related protein that is involved in beta 1 integrin-mediated signaling in lymphocytes. J Exp Med 184, 1365–1375 (1996).

39. Y. Yu, N. C. Fay, A. A. Smoligovets, H. J. Wu, J. T. Groves, Myosin IIA modulates T cell receptor transport and CasL phosphorylation during early immunological synapse formation. PLoS One 7, e30704 (2012).

40. L. C. Santos et al., Actin polymerization-dependent activation of Cas-L promotes immunological synapse stability. Immunol Cell Biol 94, 981–993 (2016).

41. D. Jevremovic et al., Cutting edge: a role for the adaptor protein LAT in human NK cell-mediated cytotoxicity. J Immunol 162, 2453–2456 (1999).

42. S. E. Degn, P. Tolar, Towards a unifying model for B-cell receptor triggering. Nat Rev Immunol 25, 77–91 (2025).

43. J. Hellmeier et al., DNA origami demonstrate the unique stimulatory power of single pMHCs as T cell antigens. Proc Natl Acad Sci U S A 118, (2021).

44. G. Voisinne et al., Kinetic proofreading through the multi-step activation of the ZAP70 kinase underlies early T cell ligand discrimination. Nat Immunol 23, 1355–1364 (2022).

45. Q. Xiao, C. K. McAtee, X. Su, Phase separation in immune signalling. Nat Rev Immunol 22, 188–199 (2022).

46. A. H. Courtney et al., CD45 functions as a signaling gatekeeper in T cells. Sci Signal 12, (2019).

47. J. M. Brockman, K. Salaita, Mechanical Proofreading: A General Mechanism to Enhance the Fidelity of Information Transfer Between Cells. Front Phys 7, (2019).

48. S. I. Roper et al., B cells extract antigens at Arp2/3-generated actin foci interspersed with linear filaments. Elife 8, (2019).

49. A. F. Carisey, E. M. Mace, M. B. Saeed, D. M. Davis, J. S. Orange, Nanoscale Dynamism of Actin Enables Secretory Function in Cytolytic Cells. Curr Biol 28, 489–+ (2018).

50. M. Fritzsche et al., Cytoskeletal actin dynamics shape a ramifying actin network underpinning immunological synapse formation. Sci Adv 3, e1603032 (2017).

51. S. Kumari et al., Cytoskeletal tension actively sustains the migratory T-cell synaptic contact. EMBO J 39, e102783 (2020).

52. H. Colin-York, S. Kumari, L. Barbieri, L. Cords, M. Fritzsche, Distinct actin cytoskeleton behaviour in primary and immortalised T-cells. J Cell Sci 133, (2019).

53. P. Bruhns et al., Specificity and affinity of human Fcgamma receptors and their polymorphic variants for human IgG subclasses. Blood 113, 3716–3725 (2009).

54. J. Wu, H. K. Mishra, B. Walcheck, Role of ADAM17 as a regulatory checkpoint of CD16A in NK cells and as a potential target for cancer immunotherapy. J Leukoc Biol 105, 1297–1303 (2019).

55. P. G. Kremer, A. W. Barb, The weaker-binding Fc gamma receptor IIIa F158 allotype retains sensitivity to N-glycan composition and exhibits a destabilized antibody-binding interface. J Biol Chem 298, 102329 (2022).

56. X. Zhao et al., Tuning T cell receptor sensitivity through catch bond engineering. Science 376, eabl5282 (2022).

57. Y. Feng, E. L. Reinherz, M. J. Lang, alphabeta T Cell Receptor Mechanosensing Forces out Serial Engagement. Trends Immunol 39, 596–609 (2018).

58. I. Wilkinson et al., Fc-engineered antibodies with immune effector functions completely abolished. PLoS One 16, e0260954 (2021).

59. S. Bournazos, A. Gupta, J. V. Ravetch, The role of IgG Fc receptors in antibody-dependent enhancement. Nat Rev Immunol 20, 633–643 (2020).

60. I. Gudelj, G. Lauc, M. Pezer, Immunoglobulin G glycosylation in aging and diseases. Cell Immunol 333, 65–79 (2018).

61. M. Steblyanko, N. Anikeeva, K. S. Campbell, J. H. Keen, Y. Sykulev, Integrins Influence the Size and Dynamics of Signaling Microclusters in a Pyk2-dependent Manner. J Biol Chem 290, 11833–11842 (2015).

62. D. F. Liu, M. E. Peterson, E. O. Long, The Adaptor Protein Crk Controls Activation and Inhibition of Natural Killer Cells. Immunity 36, 600–611 (2012).

63. J. Gohring, L. Schrangl, G. J. Schutz, J. B. Huppa, Mechanosurveillance: Tiptoeing T Cells. Front Immunol 13, 886328 (2022).

64. L. Balagopalan, K. Raychaudhuri, L. E. Samelson, Microclusters as T Cell Signaling Hubs: Structure, Kinetics, and Regulation. Front Cell Dev Biol 8, 608530 (2020).

65. I. Smyrlaki et al., Soluble and multivalent Jag1 DNA origami nanopatterns activate Notch without pulling force. Nat Commun 15, 465 (2024).

66. P. Tolar, K. M. Spillane, Force generation in B-cell synapses: mechanisms coupling B-cell receptor binding to antigen internalization and affinity discrimination. Adv Immunol 123, 69–100 (2014).

67. Y. S. Hu et al., Quantifying T cell receptor mechanics at membrane junctions using DNA origami tension sensors. Nat Nanotechnol, (2024).

68. V. P. Y. Ma et al., The magnitude of LFA-1/ICAM-1 forces fine-tune TCR-triggered T cell activation. Sci Adv 8, (2022).

