## Supplemental information for "Antibody Fc receptor CD16a mediates natural killer cell activation via mechanotransduction of piconewton forces"

**The PDF file includes:**

Materials and Methods
Figs. S1 to S14

**Other Supplementary Materials for this manuscript include the following:**

Movies S1 to S9

### Materials and Methods

### Material list

The materials used in this study are listed below.

| Material name | Source, Cat# and details | |
|---|---|---|
| **1. Antibodies** | | |
| anti-human CD16 antibody | Biolegend | 302049 |
| biotin anti-human CD16 antibody | Biolegend | 302004 |
| Alexa Fluor 647 anti-human CD16 antibody | Biolegend | 302020 |
| Alexa Fluor 594 anti-phosphotyrosine antibody | Biolegend | 309314 |
| Alexa Fluor 647 anti-LAT phospho (Tyr171) antibody | Biolegend | 946603 |
| Alexa Fluor Plus 647 F(ab')2-goat anti-mouse IgG (H+L) cross-adsorbed secondary antibody | Thermofisher | A48289 |
| PE anti-human IgG1 Fc antibody | SouthernBiotech | 9054-09 |
| phospho-p130 Cas (Tyr165) antibody | Cell Signaling | 4015 |
| Alexa Fluor 488 anti-rabbit IgG (H+L), F(ab')2 Fragment | Cell Signaling | 4412S |
| **2. Cell culture reagents** | | |
| MEM α | Thermo Fisher | 32571036 |
| DMEM, high glucose, GlutaMAX™ Supplement | Thermo Fisher | 10566016 |
| freestyle 293 media | Thermo Fisher | 12338018 |
| RPMI 1640 | Corning | 15-040-CV |
| 1× DPBS | Corning | 21-030-CV |
| Hank's balanced salt | Sigma | H8264 |
| horse serum | Thermo Fisher | 26050070 |
| fetal bovine serum (FBS) | Thermo Fisher | 10100147 |
| 10000 U/mL penicillin and streptomycin (P/S) | Thermo Fisher | 15140122 |
| 100× glutamax | Thermo Fisher | 35050061 |
| Antibiotic-Antimycotic (100X) | Thermo Fisher | 15240062 |
| 1000× $\beta$- mercaptoethanol | Thermo Fisher | 21985023 |
| 100× NEAA | Thermo Fisher | 11140050 |
| folic acid | Sigma | F8758-5G |
| myo-Inositol | Sigma | I7508-50G |
| NK Cell Isolation Kit, human | Miltenyi | 130-092-657 |
| **3. Biological reagents** | | |
| Competent Cells - DH5α | Zymo Research | T300 |
| NK92 | Dr. Oscar A. Aguilar (UCSF) | |
| NK92 CD16a | This study | |
| NK92 CD16a reporter | This study | |
| NK92 CD16a lifeact | This study | |
| K562 | ATCC | CCL-243 |
| K562 IgG1 Fc | This study | |
| K562 IgG1 Fc dimer | This study | |
| Jurkat TCR knockout | Dr. Mark Davis (Stanford) | |
| Jurkat TCR knockout NFAT reporter | This study | |
| Jurkat TCR knockout NFAT reporter CD16a | This study | |
| Lenti-X 293T cell line | Takara Bio | 632180 |
| 293TB-15 | NIH tetramer facility at Emory | |
| 293TB-15 Fc hexamer cells | This study | |

| | | |
|---|---|---|
| Human peripheral mononuclear cells (PBMCs) | Stanford Blood Center | |
| **4. Chemicals** | | |
| Blocker BSA (10%) in PBS | Thermo Fisher | 37525 |
| NucBlue Live ReadyProbes Reagent | Thermo Fisher | R37605 |
| Poly-D-Lysine | Thermo Fisher | A3890401 |
| (3-Aminopropyl)triethoxysilane | Sigma | 440140-100ML |
| Pierce Sulfo-NHS-Acetate | Thermo Fisher | 26777 |
| EZ-Link™ NHS-Biotin | Thermo Fisher | 20217 |
| Azidobutyric acid NHS ester | Lumiprobe | 63720 |
| Coverslips for sticky-Slides | ibidi | 10812 |
| sticky-Slide 18 Well | ibidi | T100B |
| Cell Activation Cocktail (without Brefeldin A) | Biolegend | 423302 |
| CK666 | Cayman Chemical | 29038 |
| Latrunculin B | Cayman Chemical | 10010631 |
| cholesterol | Thermo Fisher | A11470.18 |
| methyl-beta-cyclodextrin (m$\beta$CD) | Thermo Fisher | J66847.06 |
| Filipin III | Cayman Chemical | 70440 |
| Opti-MEM | Thermo Fisher | 31985062 |
| Fugene HD transfection reagent | Promega | E231A |
| PEG-it Virus Precipitation Solution | System Biosciences | LV810A-1 |
| Amicon® Ultra Centrifugal Filter, 10 kDa MWCO | Sigma | UFC501024 |
| SiR-actin | Cytoskeleton | CY-SC001 |
| Particles, 0.5% w/v, 5 µm | Spherotech | SVP-50-5 |
| protein A magnetic resin | Genscript | L00273 |
| **5. Recombinant proteins** | | |
| Recombinant Human IL-2 Protein | R&D systems | 202-IL-010/CF |
| Fc hexamer | expressed in house,<br>ref: J Immunol, 2018, 200 (8): 2542–2553 | |
| biotinylated human IgG1 Fc (C103S), avi tag, his tag | Acro Biosystems | IG1-H82E9-25ug |
| **6. Recombinant DNA** | | |
| Codon optimized human CD16a | IDT | |
| H2B-TagBFP2 | existing plasmid in lab | |
| ERKKTR-miRFP670nano3 | IDT/existing plasmid in lab | |
| GCaMP6F | IDT, ref: 10.1038/nature12354. | |
| NFAT-ZsGreen | IDT / plasmid in house | |
| Lifeact-mNeonGreen | IDT | |
| tethered IgG1 Fc | Thermofisher / plasmid in house | |
| tethered IgG1 Fc fragment X | Thermofisher / plasmid in house | |
| **7. Oligonucleotides** | | |
| 4 pN hairpin (IDT) | GTGAAATACCGCACAGATGCGTTTGTATAAATG<br>TTTTTTTCATTTATACTTTAAGAGCGCCACGTAG<br>CCCAGC | |
| 12 pN hairpin (IDT) | GTGAAATACCGCACAGATGCGTTTGGGTTAACA<br>TCTAGATTCTATTTTTAGAATCTAGATGTTAACC<br>CTTTAAGAGCGCCACGTAGCCCAGC | |
| 19 pN hairpin (IDT) | GTGAAATACCGCACAGATGCGTTTCGCCGCGGG<br>CCGGCGCGCGGTTTTCCGCGCGCCGGCCCGCGG<br>CGTTTAAGAGCGCCACGTAGCCCAGC | |
| 4 pN hairpin lock (IDT) | AAAAAACATTTATAC | |
| 12 pN hairpin lock (IDT) | AGA ATC TAG ATG TTA ACC | |

| | |
|---|---|
| 19 pN hairpin lock (IDT) | ACCGCGCGCCGGCCCGCGGCG |
| ligand strand (IDT) | /5AmMC6/CGCATCTGTGCGGTATTTCACTTT/3Bio / |
| Cy3B ligand strand (in house) | Cy3B/CGCATCTGTGCGGTATTTCACTTT/3Bio/, prepared in house, ref: 10.1073/pnas.1904034116 |
| BHQ2 anchor strand (IDT) | /5DBCON/TTTGCTGGGCTACGTGGCGCTCTT/3BH Q_2/ |
| biotin TGT top strand (IDT) | /5BiosG/CACAGCACGGAGGCACGACAC |
| 12 pN TGT bottom strand (IDT) | GTGTCGTGCCTCCGTGCTGTGTTTTT/3DBCON/ |
| 56 pN TGT bottom strand (IDT) | /5DBCON/TTTTTGTGTCGTGCCTCCGTGCTGTG |
| **6. Software and algorithms** | |
| SnapGene | 7.1.2 |
| FlowJo | 10.10.0 |
| Graphpad Prism 10 | 10.4.2 |
| CellTK2 | ref:10.1016/j.jbc.2023.104599 https://github.com/sjeknic/CellTK, https://celltk.readthedocs.io/en/latest/ |
| ImageJ2 | 2.16.0. https://imagej.net/software/fiji/downloads |
| 7. Instrument | |
| Nikon Ti-2 microscope | Nikon Ti-E microscope with temperature and $CO_2$ |
| Nikon CFI APO 100X OIL TIRF NA 1.49 WD 0.12MM | Nikon MRD01991 |
| RICM filter | Nikon 97271 |
| SCMOS camera | Andor Neo |
| Fluorescence filter set | Chroma ET 572/35x,632/60m |
| Fluorescence filter set | Chroma Quad cube, DAPI/FITC/TRITC/FAR RED |

### Method details

#### 1. Cells

NK92 cells were generously gifted by Dr. Oscar A. Aguilar (UCSF) to the Garcia lab. Human CD16a F158 was transduced into NK92 cells using lentiviral particles and the transduced cells were then sorted to establish the NK92 CD16a cell line. NK92 CD16a reporter cell line was subsequently engineered by transducing NK92 CD16a cells with lentiviral particles of H2B-TagBFP2, ERKKTR-miRFP670nano3, and GCaMP6f, followed by isolation of the transduced cells using FACS (Fluorescence-Activated Cell Sorting). NK92 CD16a lifeact cell line was established by transducing NK92 CD16a cells with lentiviral particles of lifeact-mNeonGreen and isolation of transduced cells using FACS. All NK92 cell lines were cultured in complete $\alpha$-MEM ($\alpha$-MEM supplemented with 10% FBS, 10% horse serum, 1× NEAA, 100 U/mL penicillin-streptomycin (P/S), 2 mM glutamax, 10 mM HEPES, 20 µM folic acid, 200 µM myo-inositol, 50 µM $\beta$- mercaptoethanol, and 200 U/mL IL-2) at 37 ºC with 5% $CO_2$.

Jurkat TCR knockout cells were gifted by Dr. Mark Davis (Stanford) to the Garcia lab. Jurkat TCR knockout NFAT reporter was generated by transducing Jurkat TCR knockout cells with lentiviral particles of NFAT-ZsGreen1. Jurkat TCR knockout NFAT reporter CD16a cells were generated by lentiviral transduction of human CD16a. K562 IgG1 Fc and K562 IgG1 Fc dimer cells were generated by transducing K562 cells with lentiviral particles of tethered IgG1 Fc or IgG1 Fc dimer. All Jurkat and K562 cell lines were cultured in complete RPMI1640 media (RPMI1640 supplemented with 10% FBS, 2 mM glutamax, 50 µM $\beta$- mercaptoethanol, and 100 U/mL P/S) at 37 ºC with 5% $CO_2$.

Lenti-X 293T and 293TB-15 were cultured in DMEM supplemented with 10% FBS, 2 mM glutamax, and 100 U P/S at 37 ºC with 5% $CO_2$.

Human peripheral mononuclear cells (PBMCs) were obtained from Stanford Blood Center under an Institutional Review Board approved protocol and kept in liquid $N_2$ after isolation at 10 million cells per mL. Prior to experiments, primary NK cells were purified from thawed PBMCs with MACS human NK cell isolation kit following manufacturer's instruction. Following purification, primary NK cells were rested for 2 h in complete $\alpha$-MEM at 37 ºC with 5% $CO_2$ before experiments.

#### 2. Plasmids and cloning

Gene fragments of interests were purchased from IDT or Thermofisher using gBlock or GeneArt custom synthesis services, or cloned from previously reported plasmids from the Covert Lab. CD16a, H2B-TagBFP2, ERKKTR-miRFP670nano3, GCaMP6f, NFAT-ZsGreen1, lifeact-mNeonGreen, IgG1 Fc, and Fc hexamer constructs were cloned into pHR lentiviral vector using Gibson assembly. All plasmids were sequenced and verified before expressing the genes of interest.

#### 3. Lentivirus packaging and transduction

Lenti-X 293T cells were seeded at $6.0\times10^5$ cells per well in a 6-well plate, and allowed to attach overnight. On the second day, cells were transfected with 260 ng PMD2.G, 500 ng psPax2, and 750 ng plasmid of interest using FugeneHD following manufacturer's instructions. The media containing plasmids was replaced with fresh complete DMEM after 16 h, and lentivirus was harvested 48 h later, and ready to infect $10^6$ cells by titration. Lentivirus was concentrated using PEG-it virus precipitation solution and resuspended in fresh media. For lentiviral transduction, 1 million cells were incubated in 5 mL fresh media containing lentivirus and 5 µg/mL polybrene for 24 h. Lentivirus was then removed by centrifugation, and cells were incubated in fresh media for another 48 h. Transduced cells were analyzed by flow cytometry 72 h post transduction to examine the expression of gene of interest.

#### 4. Protein expression

293TB-15 cells were transduced with lentiviral particles to stably secrete IgG1 Fc hexamer. After transduction, 293TB-15 Fc hexamer expressing cells were cultured in freestyle 293 media for five days and the media was collected for purification using protein A magnetic resin following manufacturer's protocol. The purified IgG1 Fc hexamer was desalted with Amicon filters and analyzed with native PAGE and denaturing SDS-PAGE gels.

#### 5. Cell sorting

Transduced cells were cultured and expanded for 5-7 days before FACS (Sony SH800S). Briefly, 10 to 20 million cells were spun down and resuspended in FACS buffer, and the subpopulation that expressed the genes of interest was sorted and cultured in complete media supplemented with 1× antibiotic-antimycotic for 2-3 days.

#### 6. Substrate preparation for force imaging and manipulation

DNA-functionalized substrates were prepared as described in literature (Fig. S9) (*67, 68*). Glass coverslips were washed thoroughly in ethanol and milliQ $H_2O$, and etched in 0.5 M KOH for 30 min. Coverslips were then washed 6 times with milliQ $H_2O$ and 3 times with ethanol, and functionalized with 3% APTES in ethanol. After 1 h, coverslips were washed with ethanol 6 times and baked at 80 °C for 20 min. The obtained amine coverslips are stored at -20 °C. To further functionalize the coverslips, azide NHS (10 mg/mL in DMSO) was added to amine coverslips and allowed to react for 24 h. Coverslips were then washed thoroughly with ethanol and incubated with sulfo-NHS acetate (10 mg/mL in DMSO) overnight. After incubation, the coverslips were washed with ethanol three times, dried, and attached to a sticky slide or bottomless adhesive microtiter plate. DNA tension probes were assembled by annealing Cy3B ligand strand, hairpin strand, and BHQ2 anchor strand (1: 1: 1.1) in 1× PBS at 95 °C for 5 min and slowly cooling down to 20 °C in 25 min using a thermal cycler. Sequence information for the oligonucleotides used for DNA tension probe technology can be found in Key Resources Table. Azide-functionalized wells were incubated with 20 nM DNA tension probes overnight at room temperature and washed three times with 1× PBS. The wells were then passivated with 1% BSA for 30 min. After passivation, wells were incubated in 10 μg/mL streptavidin for 30 min and washed three times. Biotin antiCD16 or biotin IgG1 Fc (10 μg/mL) were then added to tension probe functionalized wells for 30 min for ligand presentation, followed by three washes in 1× PBS. The wells were then ready for MTFM imaging experiments. For TGT experiments, 5 nM annealed 12 or 56 pN TGT duplex was added to a azide-functionalized glass well instead of DNA tension probes.

**7. Substrate preparation for imaging signaling dynamics**

For activating substrates, wells in glass-bottom 96 well plate were coated with 1 μg/mL streptavidin overnight at 4 °C and passivated with 1% BSA in 1× PBS for 1 h before immobilizing biotin-antiCD16 or biotin-IgG1 Fc. For non-activating controls, wells were coated with 100 μg/mL poly-D-lysine overnight at 4 °C. The wells were washed with 1× PBS immediately before use and kept in imaging buffer.

**8. Imaging experiments**

Microscopy imaging was performed using a Nikon Ti-E microscope with temperature and $CO_2$ control for live cell imaging. The microscope is equipped with Andor Neo SCMOS camera, and 10, 20, 40, and 100× objectives. Most of the signaling dynamics were acquired using the 20× objective and 3×3 binning in high throughput. Most of the tension data was acquired using the 100× objective and no binning. Cells were imaged in imaging buffer (Hank's balanced salts supplemented with 10 mM HEPES and 1% FBS.

**Imaging cell-bead interactions.** 50 μL of streptavidin beads were blocked with 1% BSA for 30 min, and then incubated with 10 μg/mL biotin IgG1 Fc in 50 μL 1% BSA at 4 °C for 4 h. The beads were then washed three times with 0.5 mL 1% BSA to obtain bead-bound Fc, which was subsequently resuspended in 50 μL imaging media for experiments. 10 μg/mL biotin IgG1 Fc was used for in solution Fc group, and 50 μL of BSA blocked streptavidin beads were used as empty bead group. Around $4×10^4$ NK92 CD16a reporter cells in 90 μL imaging media were seeded to each poly-D-lysine coated well in a glass bottom 96-well plate and allowed to attach at 37 °C for 20 min. As 10 μL of bead-bound Fc, in solution Fc, or empty beads were added to reporter cells, timelapse images were acquired in nucleus, $Ca^{2+}$, and ERK reporter channels with 1 min intervals. The bead-bound Fc in principle should have the same amount of IgG1 Fc as the in-solution control.

**Imaging cell-cell interactions.** Around $4×10^4$ NK92 CD16a reporter cells were seeded in a well coated with poly-D-lysine and allowed to attach for 20 min at 37 °C. As K562 or K562 IgG1 Fc cells were added to NK92 CD16a reporter cells at 1:1 ratio, timelapse images were acquired in nucleus, $Ca^{2+}$, and ERK reporter channels with 1 min intervals. For cell stiffness manipulation, 500 mM MβCD stock solution was prepared, and 80 mM (by cholesterol) MβCD-cholesterol complex (molar ratio: 6.5/1) solution was prepared. To remove cholesterol, cells were incubated in serum-free media containing 10 mM MβCD solution at 37 °C for 30 min. To load cholesterol, cells were incubated in serum-free media containing 500 μM (by cholesterol) MβCD-cholesterol at 37°C for 2 h. Cells were washed twice with 1 mL of PBS at room temperature before adding to NK92 CD16a reporter cells. In parallel, cells after treatment were also stained with 0.4 μg/mL Filipin III in 1×PBS for 30 min, washed, and imaged with DAPI cube to check cholesterol content.

**Imaging cell-substrate interaction.** To track $Ca^{2+}$ and ERK signaling dynamics, around $4×10^4$ NK92 CD16a reporter cells were seeded to each activating substrate. Timelapse images were acquired with 1 min interval in reporter channels with the 20× objective. To monitor NFAT activation, Jurkat TCR knockout CD16a NFAT reporter cells were pre-stained with NucBlue Live ReadyProbes Reagent to label nucleus for cell tracking and imaged with 20 min interval. For experiment groups with drug inhibition, reporter cells were pre-treated with CK666, latrunculin B, or 0.5% DMSO (control) for 20 min at 37 °C before exposed to activating substrates. To visualize actin dynamics, around $4×10^4$ NK92

CD16a lifeact cells were seeded to activating substrates or poly-D-lysine. Timelapse images were acquired starting at t = 5 min with 10 s interval for 5 min in both F-actin and RICM channel with the 100× objective. To visualize molecular forces, NK92 CD16a cells or primary NK cells were added to MTFM substrates. Both the real-time CD16a force as well as accumulated CD16a force were imaged with the absence and presence of 1 μM lock oligonucleotide following previous report (*19*).

**Immunofluorescence staining.** Cells incubated on different substrate was fixed with 4% PFA for 10 min and gently washed with 1× PBS, followed by permeabilization with 0.1% triton x-100 in 1× PBS for 10 min. Cells were then washed again with 1× PBS, blocked with 5% BSA, and stained with primary antibody overnight at 4 °C. On the second day, cells were washed and blocked again and stained with secondary antibody, followed by microscopy imaging.

**9. Flow cytometry**

**Expression of genes of interest.** Cells were stained for CD16a expression with 5 μg/mL Alexa Fluor 647 labeled antiCD16 (clone 3G8) in FACS buffer for 30 min, and washed three times before analysis on CytoFLEX flow cytometer (Beckman).

**NFAT activation.** Cells were counted and the density was adjusted to $1\times10^6$ cells per mL. 100 μL of Jurkat TCR knockout CD16a NFAT reporter cells were mixed with 100 μL K562 IgG1 Fc or K562 IgG1 Fc dimer cells and co-cultured in round-bottom 96-well plate for 16 h at 37 °C. Cells were then stained with Alexa Fluor 647 labeled anti-CD16 in FACS buffer and analyzed on CytoFLEX for NFAT activation.

**10. Quantification and statistical analysis**

Traces of $Ca^{2+}$ and ERK signaling dynamics were extracted from timelapse images using a previously reported pipeline CellTK (Figure S2, Video S1) (*30*). Briefly, nucleus segmentation was performed with U-net and thresholding to generate a nucleus mask. The nucleus mask is further expanded to generate a cytoplasm mask. The median fluorescence intensities in cytoplasm region and nucleus were obtained by applying the mask to timelapse images and used for obtaining cytoplasma/nucleus ratio. The $Ca^{2+}$ signal was the extracted median fluorescence intensity after applying the cytoplasm mask to GCaMP6f channel.

Images of receptor forces were processed following previously reported methods (*19, 33*). Briefly, the fluorescence background of the quenched probes was subtracted from the force images for representative tension images. Quantitative analysis of forces, contact area, and phosphorylation in fixed samples was performed by quantifying fluorescence signaling within the ROIs of cells, and the ROIs were determined by the outline of fluorescence signal masks and/or the outline of contact area in RICM channel.

Quantitative analysis of actin foci was performed using Image J Trainable Weka Segmentation tool (Fig. S13). Actin foci dwell time was analyzed using Image J particle tracking analysis TrackMate tool (Video S6).

Experiments were performed in triplicates. Statistical analyses were performed in GraphPad Prism and were described in each figure legend where statistical comparisons were performed, including n and p values.

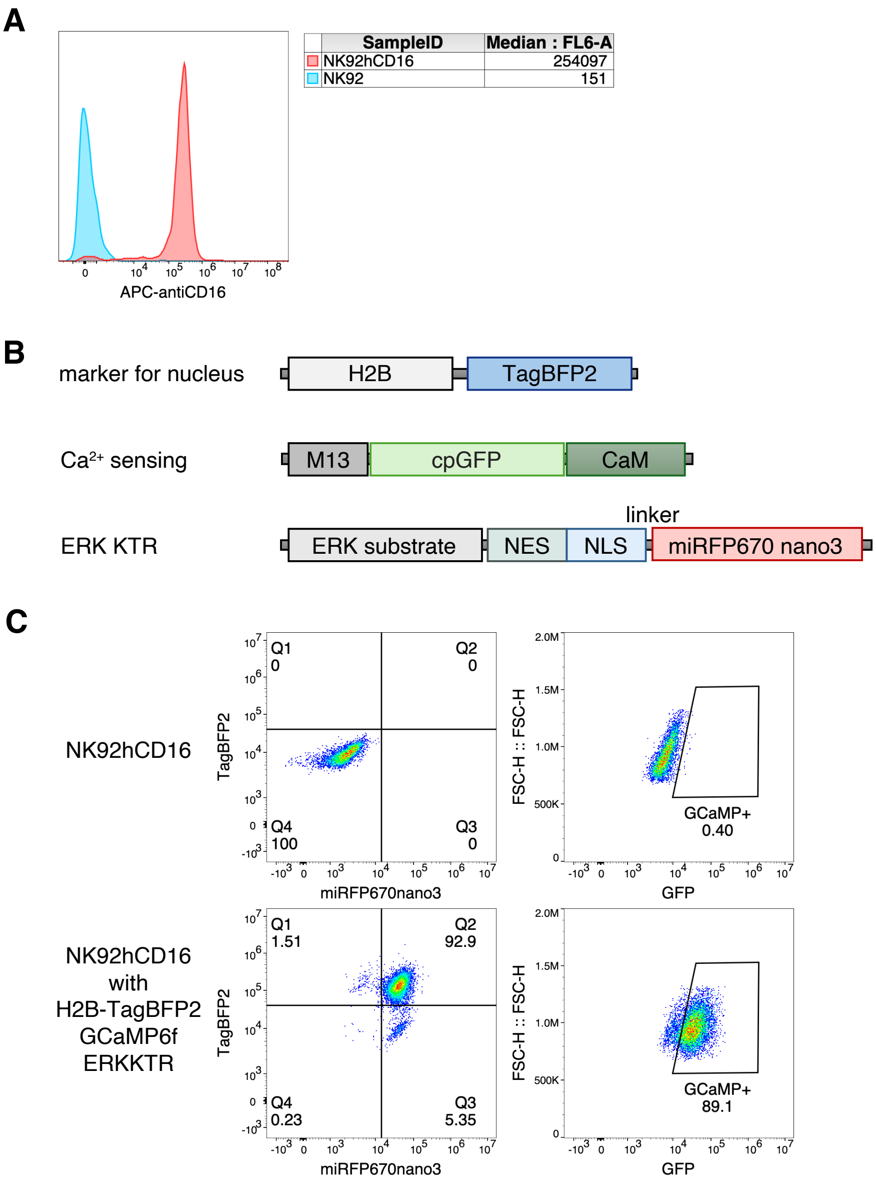


**Fig. S1. Generation of the NK92 CD16a reporter cell line.** (A) NK92 cell line was engineered to express human CD16a and the expression was confirmed by staining with APC labeled antiCD16 (clone 3G8) and analyzing with flow cytometry. (B) Constructs of fluorescent marker and reporters used to generate the NK92 CD16a reporter cell line. (C) Flow cytometry plots of NK92 CD16a cells before and after the lentiviral transduction of constructs in (B) and FACS sorting.

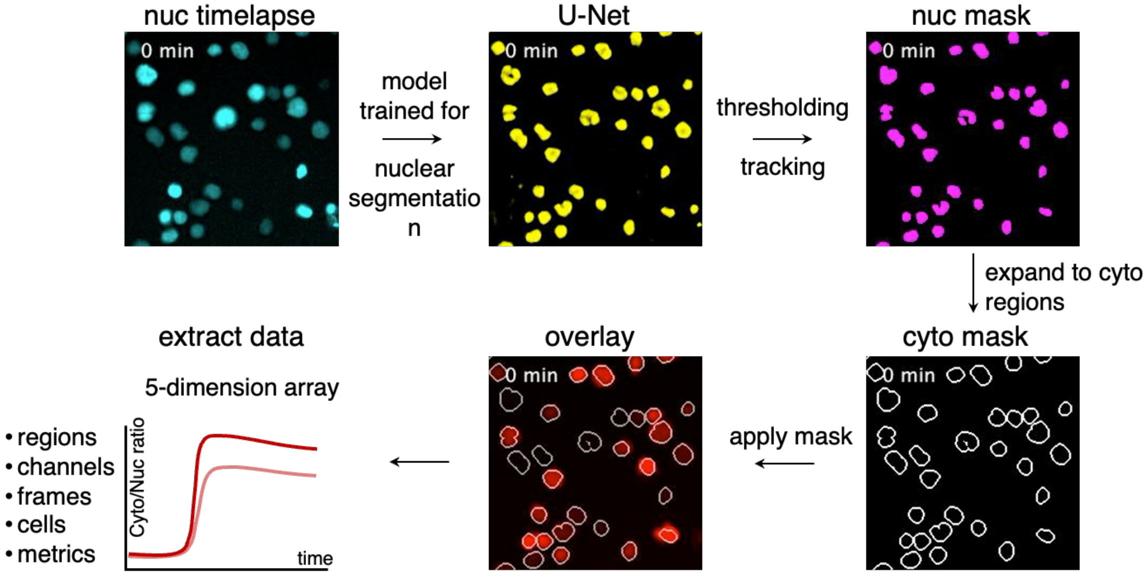


**Fig. S2. Illustration of the CellTK pipeline.** Cells were seeded on poly-D-lysine coated plates and allowed to attach for 20 min before imaging. Signaling traces were extracted using CellTK from timelapse images (*30*). Cell nucleus expressing H2B-TagBFP2 was captured using a DAPI cube and segmentation was performed using U-Net. After thresholding, a nuclear mask was generated and further expanded to cytoplasm regions to obtain the cytoplasm mask. Both masks were applied to raw images to obtain overlays and extract the corresponding data in a 5-dimention array, containing information of regions, channels, frames, cells, and metrics that users defined, such as Cyto/Nuc ratio (median fluorescent intensity of cytoplasm/median fluorescent intensity of nucleus of ERKKTR). The masks obtained were verified in overlayed timelapse videos (Video S1).

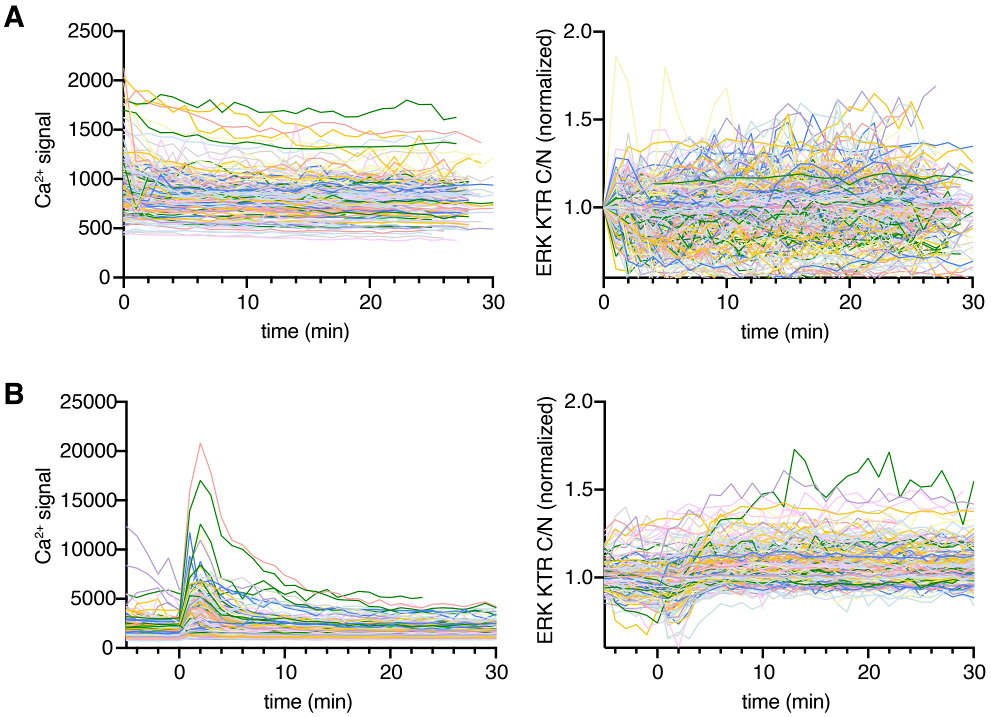


**Fig. S3. Representative extracted single-cell traces of $Ca^{2+}$ and ERK signaling activities in NK92 CD16a reporter cells.** (A) Single-cell traces of $Ca^{2+}$ and ERK signaling from ~200 cells incubated on poly-D-lysine coated glass (negative control). (B) Single-cell traces of $Ca^{2+}$ and ERK signaling from ~200 cells incubated on poly-D-lysine coated glass with PMA ionomycin (positive control).

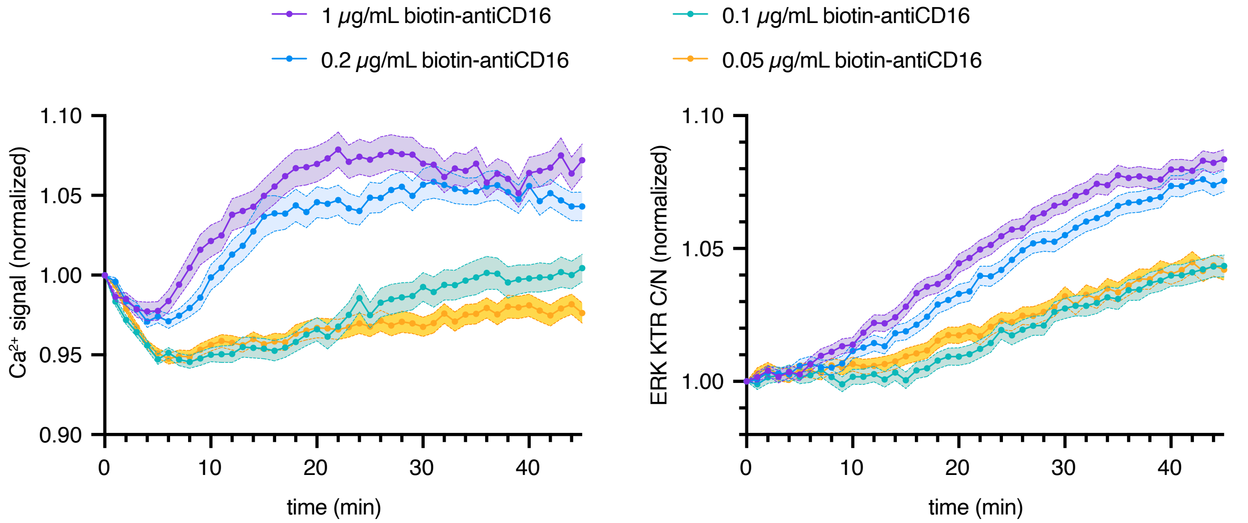


**Fig. S4. NK cell activation on glass coated with different concentrations of biotin-antiCD16.** Glass wells were coated with 1 µg/mL streptavidin overnight and passivated with 1% BSA for 1 h before immobilizing different concentrations of biotin-antiCD16. After washes, cells were seeded to each well and timelapse was immediately acquired with an interval of 1 min for 45 min at 37 °C with 5% $CO_2$. Data of each condition show mean ± SEM from >600 cells. The $Ca^{2+}$ and ERK reporters showed clear differential activation at different antiCD16 coating concentrations.

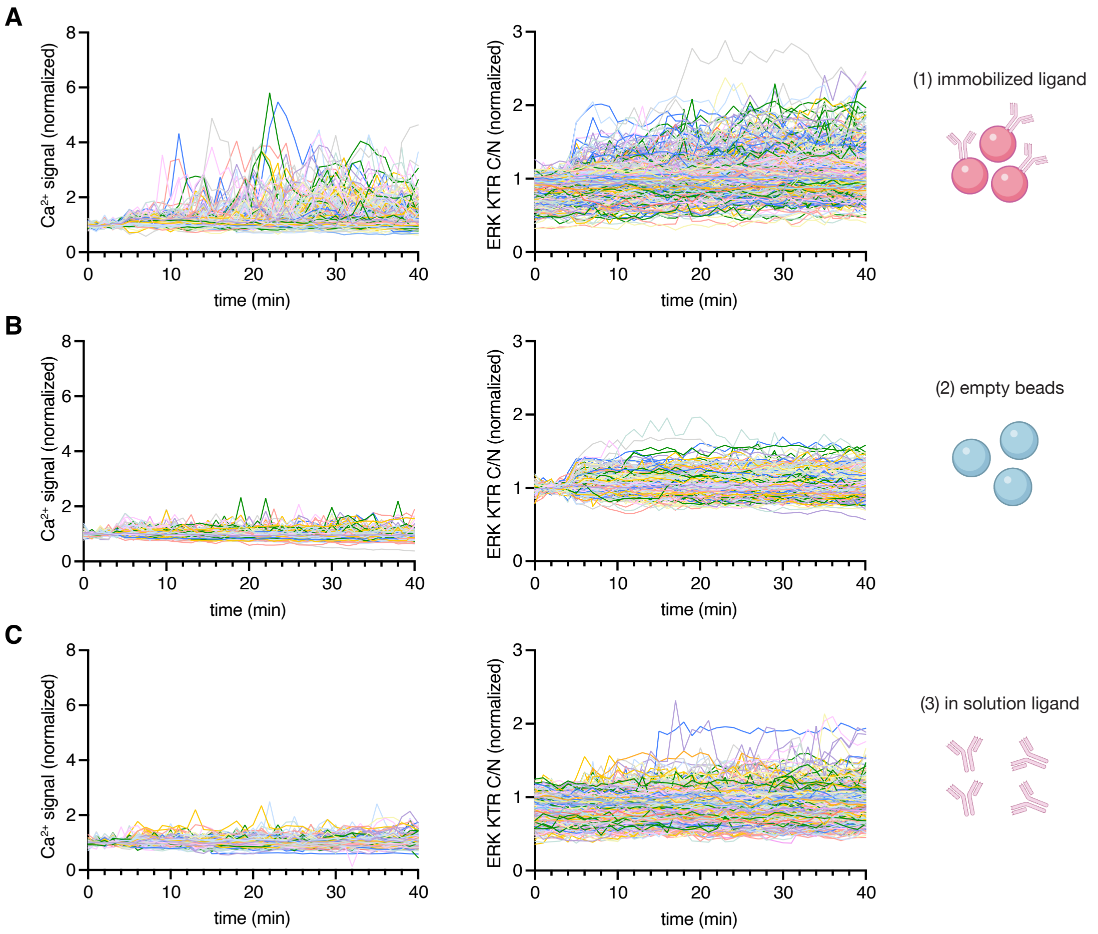


**Fig. S5. Representative single-cell traces of $Ca^{2+}$ and ERK signaling.** (A) Response from NK92 CD16a reporter cells to bead-bound Fc, n = 304 cells. (B) Response from NK92 CD16a reporter cell to empty beads, n = 363 cells. (C) Response from NK92 CD16a reporter cells to soluble Fc, n = 459 cells.

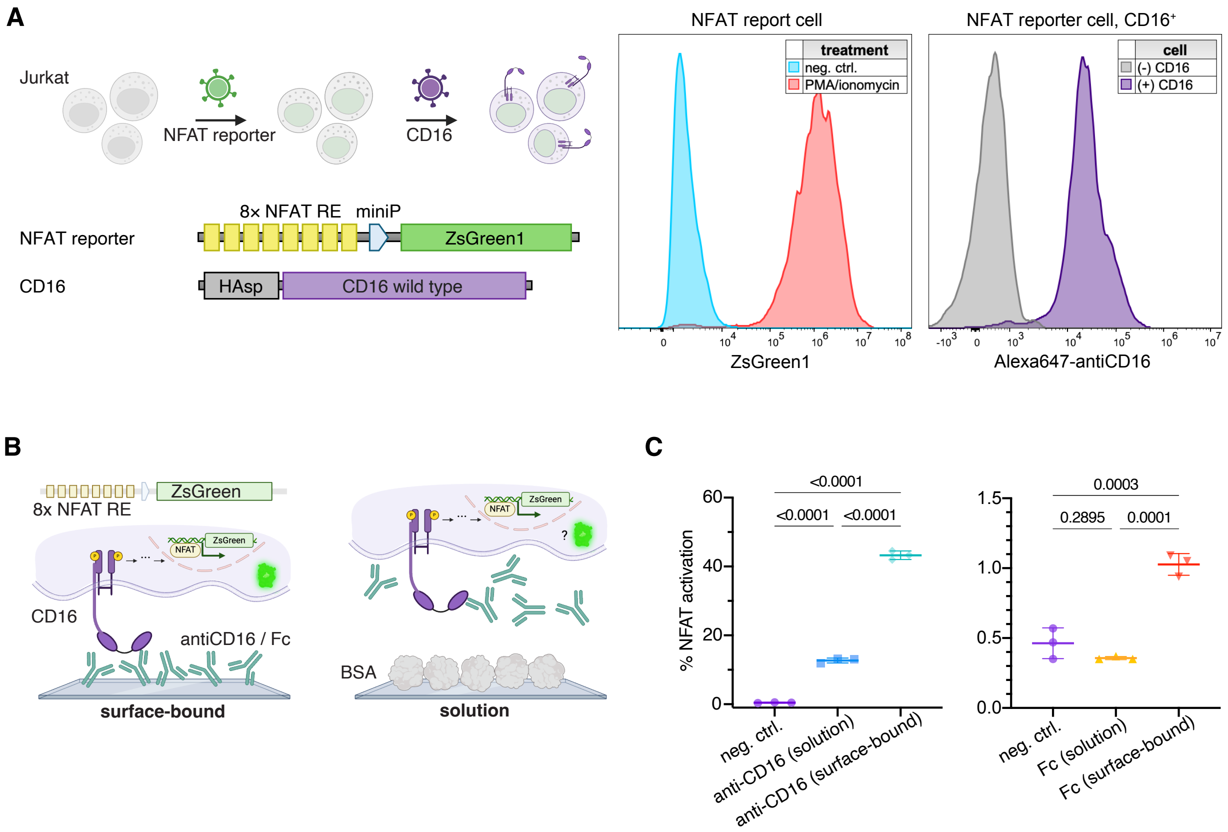


**Fig. S6. NFAT activity after Fc stimulation with an engineered Jurkat CD16a NFAT reporter cell line.** (A) Scheme and data showing engineering of the Jurkat CD16a NFAT reporter cell line. Jurkat TCR knockout cells were first transduced with lentiviral particles expressing NFAT reporter and then transduced again to express human CD16a. The sorted reporter cells after transduction respond well to PMA/ionomycin positive control and stained positive for CD16a using Alexa647-antiCD16. (B) The reporter cells were used to test the NFAT activation by soluble or surface-bound antiCD16 and IgG1 Fc. (C) Data shows that surface-bound antiCD16 results in higher % NFAT activation compared to antiCD16 in solution (mean ± SD, data was obtained from three replicates).

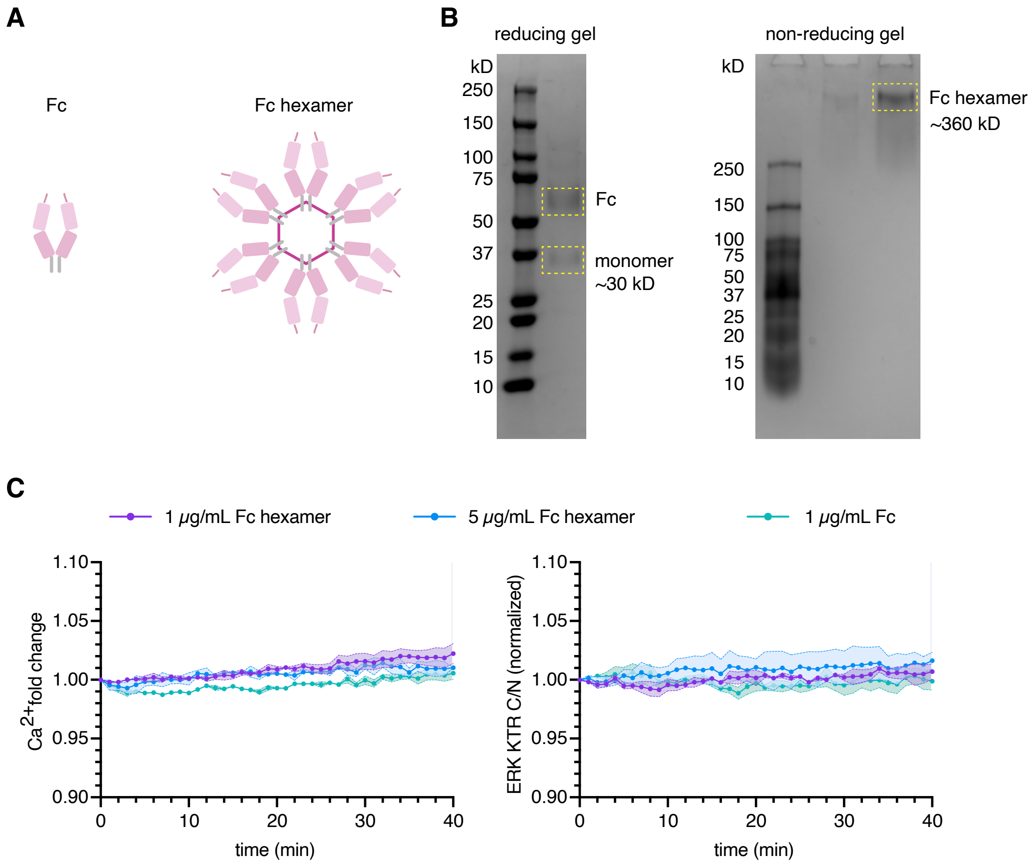


**Fig. S7. Fc hexamer expression and NK cell activation.** (A) Fc vs. Fc hexamer. (B) Reducing and non-reducing protein gels of expressed Fc hexamer after purification showing monomers at ~ 30kDa and assembled hexamers at ~ 360 kDa. (C) Representative population response of $Ca^{2+}$ and ERK signaling in NK92 CD16a reporter cells upon incubation with Fc or Fc hexamer in solution. Mean±SEM, n ~ 50 cells per condition in each replicate, data was obtained from three replicates.

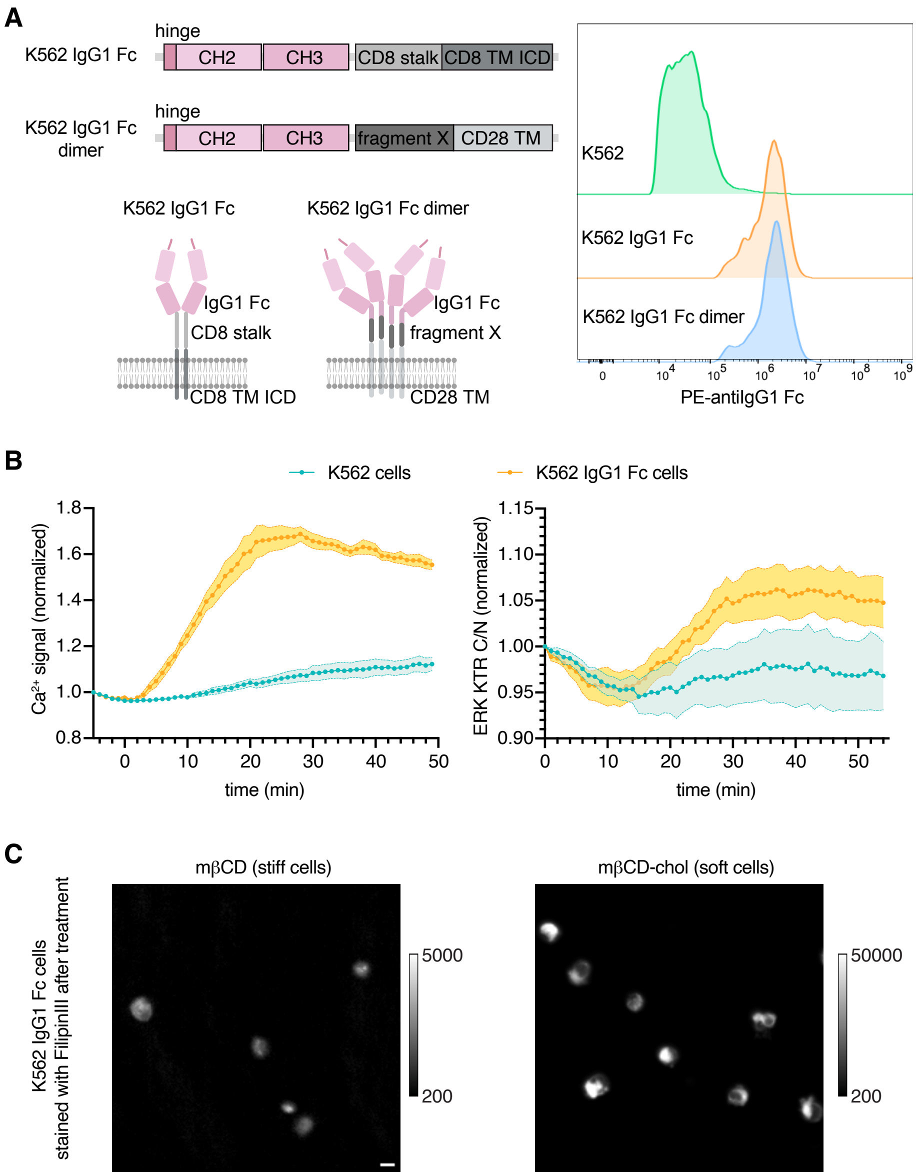


**Fig. S8. Engineering K562 IgG1 Fc target cell.** (A) Constructs of cell membrane-tethered Fc were created and cloned into a lenti vector. K562 cells were transduced, and the expression of surface Fc was confirmed by flow cytometry using a PE-antiIgG1 Fc antibody. (B) $Ca^{2+}$ and ERK response from NK92 CD16a reporter cells co-incubated with K562 or K562 IgG1 Fc cells. Data shows the mean±SEM, averaged from ~200-400 single cell traces in each run, and the experiment was performed in triplicates. (C) K562 IgG1 Fc cells were treated with mβCD or mβCD-cholesterol, stained, and imaged for cholesterol content using FilipinIII. Note the calibration bars are different. Scale bar = 10 μm.

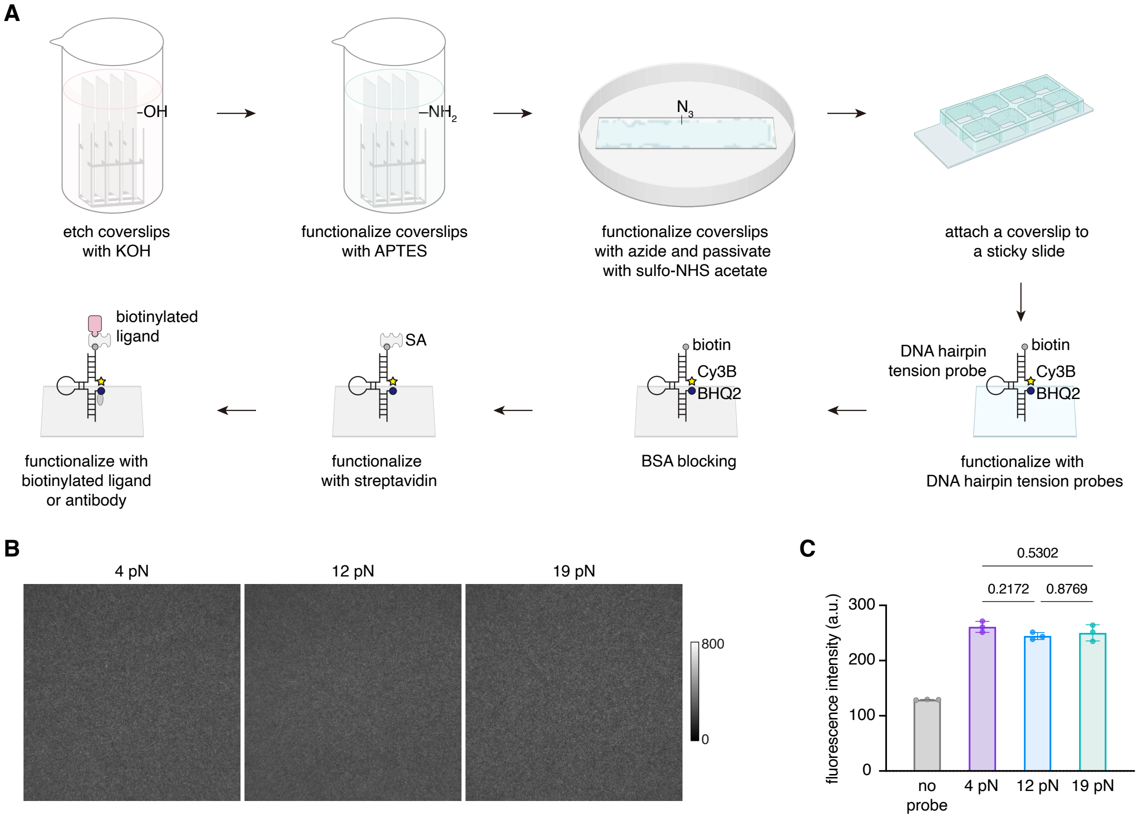


**Fig. S9. Substrate preparation and characterization.** (A) Scheme illustrating coverslips were first etched in KOH and functionalized with amines for conjugation with azide-NHS. The azide modified coverslips were than passivated and attached to a chamber. The coverslip is functionalized by DNA tension probes with DBCO modification and subsequently blocked by BSA solution. Streptavidin was added to immobilized biotinylated antibody or ligand. (B) Representative fluorescent images of DNA tension probe substrate (Cy3B signal, collected using TRITC filter setting) with $F_{1/2}$ values of 4, 12 , and 19 pN. (C) Quantitative analysis shows that the substrates of different force threshold have identical density. Measurement was obtained from 3 replicates, and 3 to 5 random positions from each replicate with 100× objective.

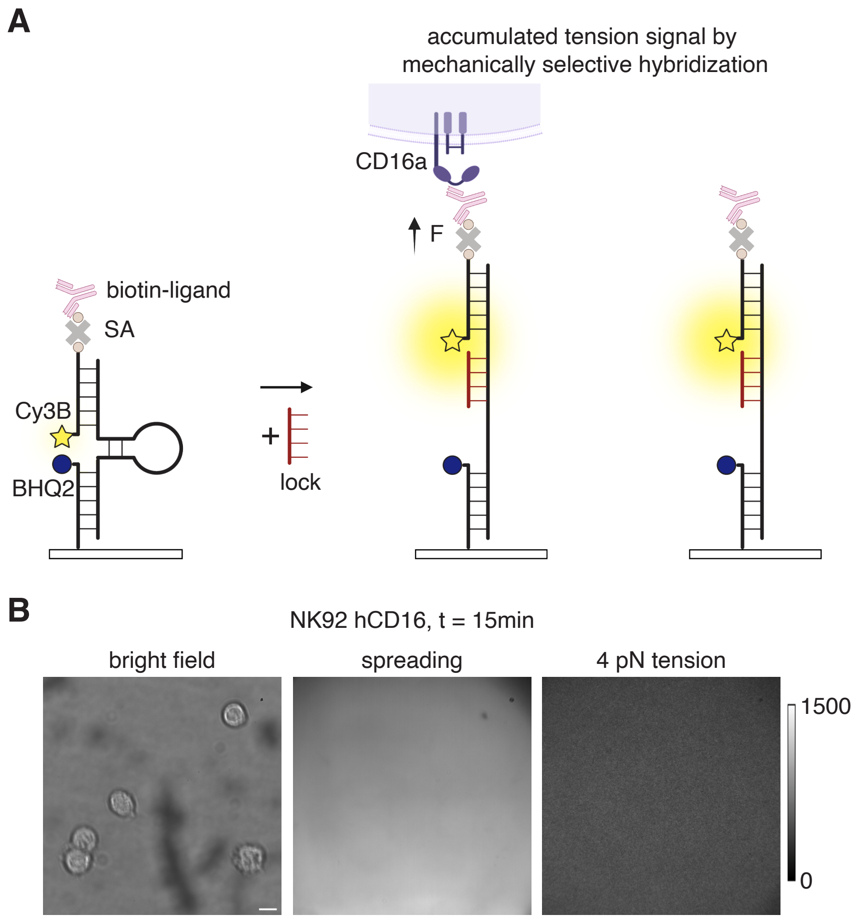


**Fig. S10. Controls of DNA tension probe measurements.** (A) Schematic showing measuring accumulated force with the lock oligonucleotide. Once lock is added, real-time tension signal is chemically accumulated through hybridization and therefore generate irreversible fluorescence signal upon mechanical unfolding. (B) NK92 CD16a cells were incubated on DNA probe substrates with streptavidin but without biotinylated ligands as the negative control group to confirm the specificity of CD16a forces shown in Fig. 2. No spreading or tension signal were observed. Scale bar = 5 µm.

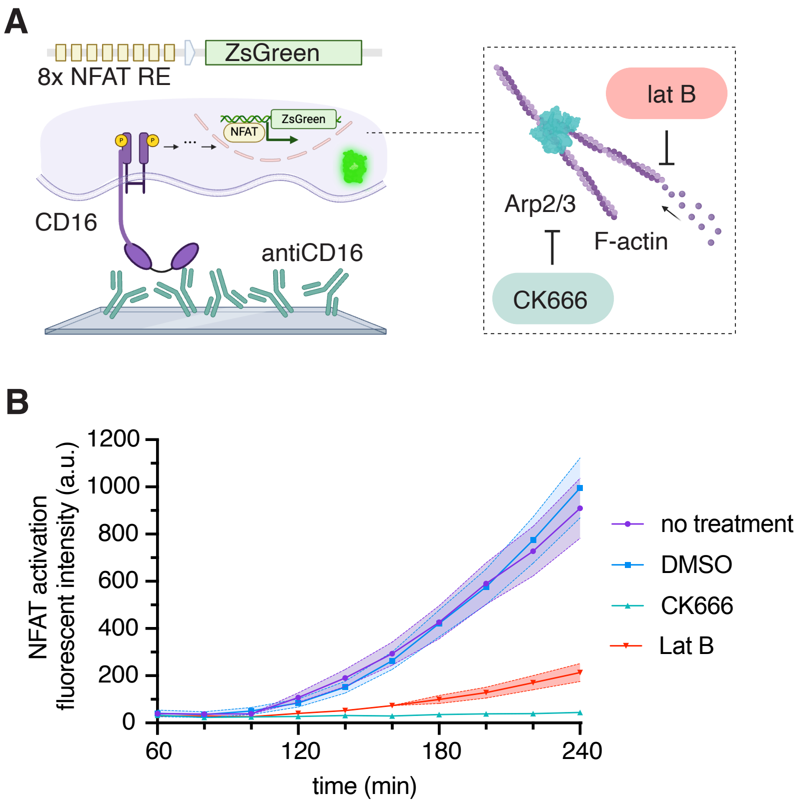


**Fig. S11. NFAT activation was inhibited by CK666 and lat B treatment in Jurkat CD16a NFAT reporter cells.** (A) Scheme showing Jurkat CD16a NFAT reporter cells were pretreated with DMSO, CK666 or lat B at 37 °C for 20 min, and then plated on antiCD16 coated wells for activation. (B) Single cells were tracked with a live nuclear stain during imaging and activation in NFAT was measured by ZsGreen1 intensity. Data show mean±SEM from three replicates, n = 691, 954, 2006, and 1653 cells for no treatment, DMSO, CK666, and lat B group respectively. Cytoskeleton inhibition significantly dampens NK activation through CD16a.

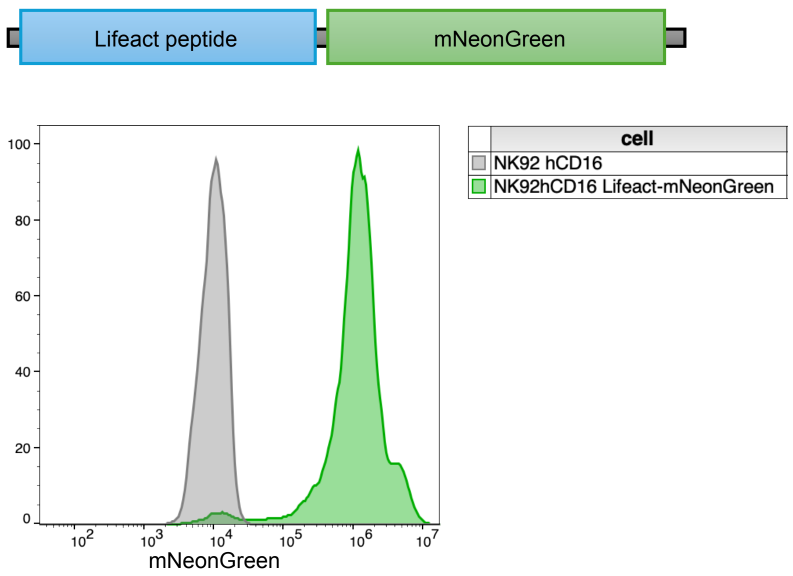


**Fig. S12. NK92 CD16a expressing F-actin marker.** NK92 CD16a cells were transduced with lentiviral particles to express the Lifeact-mNeonGreen. Lifeact is a short peptide MGVADLIKKFESISKEE and binds to filamentous actin, which can be used for imaging F-actin dynamics. The stably expressing cells were sorted by FACS and cultured for experiments where visualization of F-actin dynamics was required.

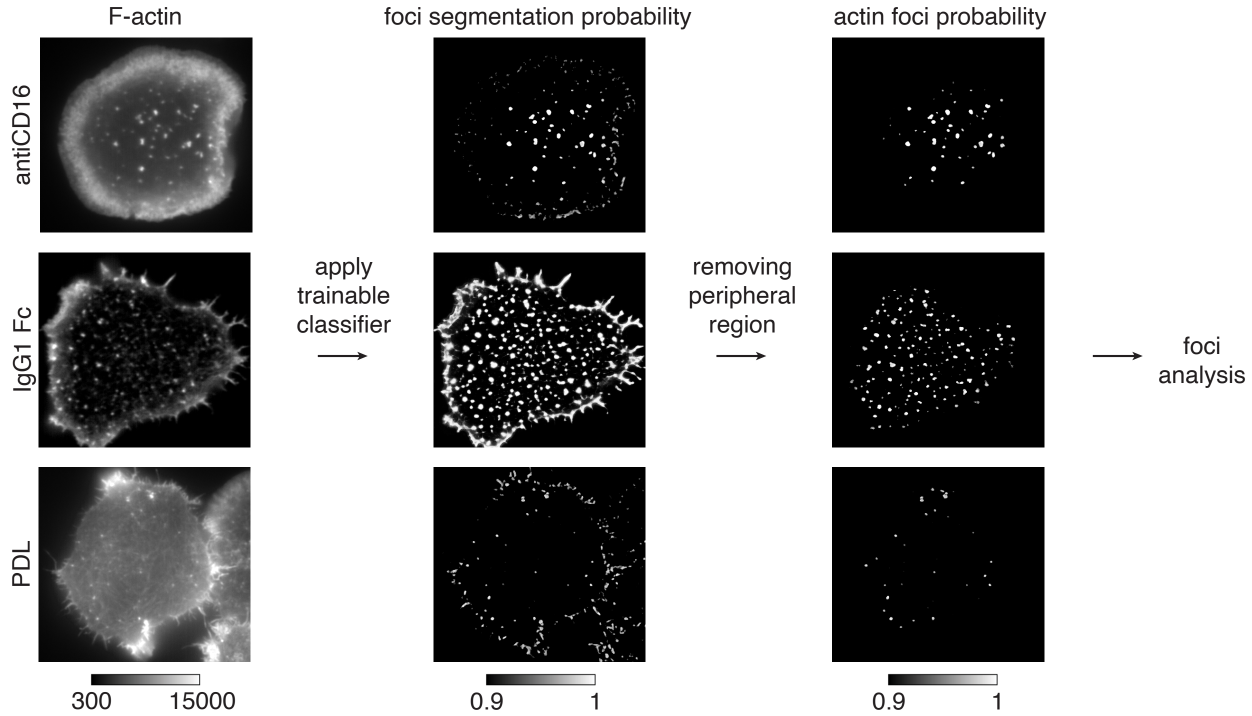


**Fig. S13. Actin foci quantification in NK92 CD16a lifeact-mNeonGreen cells.** Images of cells were collected in lifeact-mNeonGreen channel, and the actin foci were segmented using a trainable classifier. The classifier was then generated using Image J Trainable Weka Segmentation tool. The classifier was applied to images of cells to obtain segmented actin foci with > 0.9 probability for quantitative analysis.

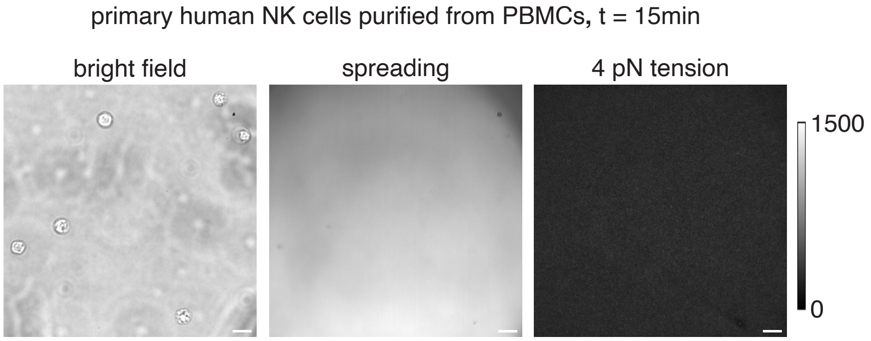


**Fig. S14. Primary human NK cells were incubated on DNA probe substrates with streptavidin but no biotinylated ligands.** Bright field images indicate the presence of the cells, but no contact or tension was observed in RICM and tension channels after 15 min of incubation. Scale bar = 5 µm.

Video S1. Single-cell signaling dynamics tracking with CellTK

Video S2. $Ca^{2+}$ and ERK signaling when NK cells interact with stiff or soft target cells

Video S3. NK92 CD16a cells transmitting forces above 4 pN against human IgG1 Fc

Video S4. Activation of NK92 CD16a reporter cells on 4, 12 , and 19 pN hairpin tension probes

Video S5. F-actin foci dynamics on different substrates. Left to right: poly-D-lysine, antiCD16, and human IgG1 Fc.

Video S6. TrackMate Capture of F-actin foci on different substrates.

Video S7. CD16a force transmission and F-actin dynamics

Video S8. Primary NK cells generating CD16a forces greater than 4 pN against IgG1 Fc

Video S9. Primary NK cells generating CD16a forces greater than 4, 12, or 19 pN against antiCD16
